# Suppression gene drive in continuous space can result in unstable persistence of both drive and wild-type alleles

**DOI:** 10.1101/769810

**Authors:** Jackson Champer, Isabel Kim, Samuel E. Champer, Andrew G. Clark, Philipp W. Messer

**Affiliations:** Department of Computational Biology, Cornell University, Ithaca, NY 14853; Department of Molecular Biology and Genetics, Cornell University, Ithaca, NY 14853

## Abstract

Rapid evolutionary processes can produce drastically different outcomes when studied in panmictic population models versus spatial models where the rate of evolution is limited by dispersal. One such process is gene drive, which allows “selfish” genetic elements to quickly spread through a population. Engineered gene drive systems are being considered as a means for suppressing disease vector populations or invasive species. While laboratory experiments and modeling in panmictic populations have shown that such drives can rapidly eliminate a population, it is not yet clear how well these results translate to natural environments where individuals inhabit a continuous landscape. Using spatially explicit simulations, we show that instead of population elimination, release of a suppression drive can result in what we term “chasing” dynamics. This describes a condition in which wild-type individuals quickly recolonize areas where the drive has locally eliminated the population. Despite the drive subsequently chasing the wild-type allele into these newly re-colonized areas, complete population suppression often fails or is substantially delayed. This delay increases the likelihood that the drive becomes lost or that resistance evolves. We systematically analyze how chasing dynamics are influenced by the type of drive, its efficiency, fitness costs, as well as ecological and demographic factors such as the maximal growth rate of the population, the migration rate, and the level of inbreeding. We find that chasing is generally more common for lower efficiency drives and in populations with low dispersal. However, we further find that some drive mechanisms are substantially more prone to chasing behavior than others. Our results demonstrate that the population dynamics of suppression gene drives are determined by a complex interplay of genetic and ecological factors, highlighting the need for realistic spatial modeling to predict the outcome of drive releases in natural populations.

## INTRODUCTION

Where evolution occurs over timescales that are short compared to the time it takes alleles to disperse across a population by migration, modeling based on the assumption of panmixia is often inadequate, and frameworks that explicitly incorporate spatial structure must be utilized to accurately predict evolutionary dynamics^1–8^.

Taking the idea of rapid evolution to an extreme, CRISPR gene drives can, in principle, spread through a population in just a few generations due to super-Mendelian inheritance^9–14^. Such drives could provide new approaches in the fight against vector-borne diseases and invasive species^9–11,13^. For example, a modification-type drive could be engineered to spread an anti-malaria gene through a mosquito population, replacing wild-type individuals with drive carriers that cannot transmit the disease. Alternatively, a suppression-type drive could be used to eliminate a disease vector or an invasive pest species. That such ideas are no longer the realm of science fiction was highlighted by a recent study that successfully eliminated cage populations of *Anopheles gambiae* with an engineered CRISPR suppression drive^15^. However, it remains unclear how such drives would perform in natural populations that are spatially structured, where the spread of alleles is constrained by the movement of individuals.

Panmictic population models have been useful in identifying optimal gene drive parameters and studying the effects of phenomena such as the evolution of resistance^16–19^. Yet it has also become clear that to accurately understand the full range of outcomes of a gene drive release, spatial factors must be explicitly considered^20–25^. Panmictic models typically predict, for instance, that a suppression drive will either successfully eliminate a population or be quickly lost. By contrast, in models that consider space, a drive might initially suppresses a population locally, but then eliminate itself before spreading to wild-type individuals in surrounding areas^22^. Spatial structure can also lead to coexistence of drive and wild-type alleles where panmictic models would predict that one would always outcompete the other quickly. For example, in a mosquito-specific simulation of a malaria-endemic nation represented by a network of linked panmictic populations, “metapopulation dynamics” were frequently observed, wherein a drive invasion resulted in local population elimination, which was then followed by recolonization by wild-type individuals^24^. Another study showed that an equilibrium can exist between empty space, regions with only wild-type alleles, and regions also containing drive alleles^26^.

The observation of coexistence of drive and wild-type alleles in spatial suppression drive models evokes a spatial game of rock-paper-scissors, wherein the coexistence of three cyclically dominant classes is facilitated^27–30^. For a suppression gene drive, these classes are drive alleles, wild-type alleles, and empty space. Drive alleles replace wild-type alleles (through conversion), wild-type alleles replace empty space (through colonization), and empty space replaces drive alleles (through drive-induced suppression). Thus, we may potentially expect suppression gene drives to exhibit complex dynamical patterns in spatial populations similar to those that have been observed in other natural systems with spatial rock-paper-scissors dynamics^28,29^.

In this study, we systematically explore how a population model with continuous space affects the dynamics and outcome of suppression gene drives. We will show that a “chasing” phenomenon arises that is not observed in panmictic population models. Chasing describes spatio-temporal dynamics similar to rock-paper-scissors that can lead to long-term coexistence between drive and wild-type alleles. We analyze the propensity of different types of suppression drives to result in chasing dynamics and show how this phenomenon depends on the ecological and demographic parameters of the population.

## METHODS

### Suppression drive strategies

We studied four gene drive strategies for population suppression, each of which is capable of rapid population elimination in a panmictic model:

1. <u>*Female fertility homing drive*</u>. This is a CRISPR/Cas9 homing drive that cleaves the wild-type allele of a heterozygote in the germline and then copies itself into that location by homology-directed-repair. We assume that the drive allele is placed inside a haplosufficient female fertility gene, inactivating the gene by its presence. Drive homozygous females (or females with any combination of drive alleles and resistance alleles that also disrupt the target gene) are sterile. As the drive increases in frequency, an accumulation of sterile females causes the population to collapse. If cleavage repair takes place by end-joining rather than homology-directed-repair (in the germline as an alternative to homology-directed repair or in the embryo due to maternally deposited Cas9), guide RNA (gRNA) target sites are often mutated. This typically creates a nonfunctional version of the target gene (called an “r2” resistance allele) that can no longer be cleaved. Such r2 resistance alleles do not typically pose major issues for this drive, since they usually don’t prevent population elimination, even though they reduce overall drive efficiency. A more severe problem is posed by “r1” resistance alleles, mutations that prevent targeting by gRNAs and preserve target gene function. However the formation rate of r1 alleles can be reduced by using multiple gRNAs^31^ or a highly-conserved target site that cannot tolerate mutations^15^. In a recent experiment, a female fertility homing drive like the one we model here was successful in rapidly eliminating small cage populations of *Anopheles gambiae*^15^.
2. <u>*Both-sex fertility homing drive*</u>. This drive is similar to the female fertility homing drive except that the drive resides in a gene that is required for both female and male fertility. Such a drive may be easier to engineer in some species if it uses a more common type of target gene. We also considered a variant of this drive that targets a recessive lethal gene with lethality at the embryo stage, another common type of gene.
3. <u>*Driving Y*</u>. The third suppression system is a Driving Y chromosome, which involves inserting the drive on the Y chromosome. In the “X-shredder” variant we consider here, the drive cleaves sites located on the X chromosome during meiosis^22,24^. With a high cleavage rate, most X chromosomes in the germline of a drive-carrying male are destroyed, resulting in most viable sperm containing the Y chromosome. As the drive spreads, the sex-ratio becomes increasingly male-biased until the number of females is so low that the population collapses. While this drive is well-studied theoretically, it has proven difficult to engineer due to low expression rates of transgenes on the Y chromosome (though autosomal X-shredders have been successfully developed^32,33^).
4. <u>*Toxin-Antidote Dominant Sperm (TADS) suppression drive*</u>. This drive does not spread by homing but relies on toxin-antidote principles to increase in frequency. Here, the “toxin” is the Cas9 with gRNAs targeting an essential spermatogenesis gene for disruption, and the “antidote” is a recoded, cleavage-resistant copy of this gene that is included in the drive allele^34^. The target gene is specifically expressed after meiosis I in males, with this expression being critical for spermatogenesis^34^. The drive allele resides in a recessive male fertility gene, disrupting the gene with its presence and causing sterility in male drive homozygotes. When Cas9 cleavage is repaired (by either end-joining or by homology-directed repair with a disrupted allele as a template), this typically creates a loss-of-function mutation. Sperm exposed to the toxin will thus not mature unless they are “rescued” by the drive. The drive spreads mainly through male heterozygotes, and the population declines as homozygous males accumulate. Maternal Cas9 activity helps this drive by creating more disrupted target alleles that will be removed from the population. Like the Driving Y system, a TADS suppression drive may prove difficult to engineer, in this case because it requires a highly specific target gene.

### Simulation model

To study the expected population dynamics of these four suppression drives, we created a simulation model of a sexually reproducing diploid population evolving over discrete, non-overlapping generations. All simulations were implemented in the forward-in-time genetic simulation framework SLiM, version 3.2.1^35^.

Gene drives were modeled to operate during offspring generating according to the following rules: First, an offspring randomly receives an allele from each parent. For the female fertility homing drive and both-sex fertility homing drive, a wild-type allele obtained from a parent that also has a drive allele is converted to a resistance allele at a rate specified by 1 – *drive efficiency* (we set *drive efficiency* = 0.95 as default, assuming an effective Cas9 promoter^15^). These resistance alleles disrupt the target gene unless otherwise specified. If a target site remains wild-type, it is converted into a drive allele at a rate equal to the *homology-directed-repair success rate*, set to 0.99 in all cases in this study. These processes actually take place in the germline prior to allele inheritance. The allele remains wild-type if neither of these events occur. However, if the mother possessed a drive allele and the offspring has remaining wild-type target sites (regardless of which parent they came from), they may be converted into resistance alleles as a result of maternal Cas9 activity in the embryo. This occurs at a rate specified by the *embryo resistance rate*, which was set to 0.05 for all homing drives, a rate corresponding to a good Cas9 promoter^15^.

For the Driving Y, a wild-type allele represents an intact X chromosome. If an offspring initially receives an undisrupted X chromosome from a father who also carried a driving Y chromosome, then X-shredding occurs at a rate specified by the *drive efficiency* (set to 0.99 by default for this drive). If the X chromosome was shredded, then the child’s genotype is redrawn, since a sperm containing a shredded X would not be viable. If the offspring ultimately receives a drive allele, then it is male. If it receives an intact wild-type X chromosome from their father, it is female.

For the TADS suppression drive, offspring receive two alleles from each parent. The first represents the target spermatogenesis gene, and the second represents the drive site (which is either a drive allele or a functional male fertility gene). If an offspring inherits a wild-type target gene from a parent who also had a drive allele, then this target gene is disrupted (with this actually taking place in the germline) at a rate equal to the *drive efficiency* (set to 0.99 by default for this drive). If an offspring has a disrupted target gene from a drive-carrying father yet lacks a paternal drive allele, then its genotype is redrawn, since such sperm would not have matured. For this system, additional Cas9 cleavage in the embryos of drive-carrying mothers does not hamper drive performance and is in fact desirable. Thus, we assume a Cas9 promoter that results in high cleavage activity in both the germline and the embryo, with the *embryo cut rate* set at 95% of the *drive efficiency*.

### Panmictic population model

We first implemented these simulations in a panmictic population, assuming the following life cycle: Generations begin with reproduction. Each non-sterile female randomly samples a male from the population. Once sampled, a male’s probability of becoming the mate is equal to 0.5, multiplied by his genotype-based fitness. This method was used to simplify computational requirements compared to assigning each male a fitness value at the outset, while assuring that the probability of being chosen remains proportional to each male’s fitness. Fitness costs are assumed to be multiplicative, such that drive homozygotes (or drive-carrier males for the Driving Y) have a fitness equal to the *drive fitness* (set to 0.95 by default), while drive heterozygotes have fitness equal to the square root of this value. When investigating inbreeding effects, we scale this probability such that male siblings become more or less likely to be selected than unrelated males. Each female makes up to 20 attempts to find a suitable mate. If she fails to find one, or if she ultimately selects a sterile male (an individual without any wild-type or functional r1 resistance alleles in the both-sex sterile homing drive, drive homozygotes for the TADS suppression drive, or a male with two disrupted TADS target genes without any drive alleles), she will have no offspring. This mating behavior is representative of mosquito populations where females typically only mate once.

In the panmictic population model, once a female has chosen a mate, we scale her fecundity to *ω_i_*′ = *ω_i_* * *β* / [(*β* − 1) *N* / *K* + 1], where *N* is the population size, *β* specifies the *low-density growth rate* of the population, *K* the *carrying capacity*, and *ω_i_* is the fecundity based on genotype. We then draw the number of her offspring from a binomial distribution with *n* = 50 and *p* = *ω_i_*′ / 25. This density dependence produces logistic growth dynamics and should push the population toward carrying capacity (in the absence of a suppression drive). If the population size is near capacity, fecundity should tend towards *ω_i_′* ≈ 1, resulting in two offspring on average. However, if the population size is lower than capacity, females will tend to produce more than two offspring. We chose a value of *β* = 6 as the default low-density growth rate, based on estimates of the *Anopheles* rate that ranges between 2 and 12^16^.

Simulations were initialized by allowing an initial wild-type population of *K* = 50000 individuals to evolve over 10 generations to reach an equilibrium. Then, drive-carriers, heterozygous for a drive and wild-type allele, were released at a frequency of 1% of the total population. For the driving Y, only male drive carriers were released, while releases comprised males and females at equal proportions for all other drives.

### Two-dimensional spatial model

We extended our model into continuous space by tracking every individual’s position across a 1×1 square landscape. The generation cycle begins with reproduction. However, rather than sampling any male from the population as a potential mate, females are now restricted to sampling from within a radius specified by the *migration rate*, which we set to 0.04 by default. If the female cannot find a suitable mate within this radius, then she cannot reproduce. In this model, we assume that density regulation is local. In particular, we define a *carrying density*, *ρ*_*k*_ = *K* / *total area*, and then compare this value to the *local density*, *ρ_i_*, around a female, defined as the density of individuals within a circle of radius 0.01 around her. This interaction function is inspired by mosquito larval competition, in which all larvae in the same pool of water compete, but larvae in adjacent pools do not compete with one other.

The fitness of the female is then scaled to *ω*_*i*_′ = *ω*_*i*_ * *β* / [(*β* − 1) *ρ*_*i*_ / *ρ*_*k*_ + 1], where *β* represents the low-density growth rate of the population. This model allows females in a less densely populated area to have higher fitness due to less competition for resources. Finally, the number of offspring is drawn from a binomial distribution with *n* = 50 and *p* = *ω*_*i*_′ / 25, such that females in areas of very low-density will have, on average, *2β* offspring. Such density-dependence could represent reduced competition at various stages of the life cycle, depending on the biological system. For mosquitoes, the benefits of reduced competition are primarily at the larval phase.

Offspring genotypes are obtained according to the same suppression drive mechanisms as in the panmictic model. Once an offspring’s genotype is determined, the individual is displaced from the position of its mother in a random direction by a random distance drawn from a normal distribution with mean zero and standard deviation equal to the *migration rate*. Coordinates that fall outside of the arena are redrawn until they fall within the boundaries.

Our spatial simulations were initialized by scattering a population of *K* = 50000 wild-type individuals (the *population capacity*, which is allowed to vary only in Figure S8) uniformly across the landscape. We then allowed the population to equilibrate for 10 generations before releasing drive heterozygotes (or drive-carrying males in the driving Y) at 1% of the total population frequency in a central circle of radius 0.01.

### Analyses of simulation outcomes

In our simulations, we recorded allelic frequencies, population size, and the frequency of sterile males and females at the end of every generation. We allowed each simulation to run for 1000 generations, but stopped the simulation earlier if the population was eliminated, the drive allele was lost while wild-type individuals were still present, r1 resistance alleles evolved and prevented the spread of the drive (by reaching 10% total frequency), or the drive fixed but the population was not eliminated within 10 generations (this only occurred when an inefficient Driving Y fixed in the population and was considered to be an “equilibrium” outcome).

To quantify the degree of clustering in the spatial population model at any given time step, we calculated Green’s coefficient^36^. We first divided the landscape into an 8×8 grid of equal-sized square cells, and counted the number of wild-type homozygote individuals, *n*_*i*_, in each cell. Green’s coefficient is then defined as *G* = (*s*^2^/*n*−1)/(*N*−1), where *n* and *s*^2^ are the mean and variance, respectively, of the individual cell counts, and *N* denotes the total population size. If individuals are randomly distributed over the space according to a Poisson distribution, then *n* and *s*^2^ should be equal, yielding *G* = 0. Clustering of individuals in space leads to *s*^2^ > *n*, and thus *G* > 0. The maximum value is *G* = 1, specifying a scenario in which all individuals are located within a single cell. Values of *G* between 0 and 1 allow quantification of the degree of spatial clustering in the population. Note that we only count wild-type homozygotes in our estimation of *G*, as we found this to produce a larger dynamic range than when all individuals were included.

Based on the inferred time-series *G*(*t*) and *N*(*t*) in a simulation run, we developed an ad-hoc procedure to decipher if and when chasing occurred for instances where the drive did not fix and reach an equilibrium population. The drive initially clears the population radially from the center of the landscape, causing wild-type individuals to become increasingly clustered around the edges. This results in increasing *G*(*t*) and decreasing *N*(*t*). However, when wild-type individuals escape to low-density areas and rebound, starting a chase, this pattern is reversed. We aimed to capture this scenario by identifying the first maximum in Green’s coefficient and minimum of wild-type allele count (with monotonic decrease or increase, respectively, required for three generations on either side of the extrema) after the population had declined by at least 20% from its starting size. This indicates the point when wild-type individuals start to increase again, and clustering of wild-type individuals start to decrease due to the expanding wild-type population during a chase. If there was an extremum in both, we considered a chase to have occurred. We considered the lowest generation of the two extrema to be the generation that chasing began. We tested this method by visually identifying the generation in which a chase began and comparing our visual detection to this automated test. Under a wide variety of parameters and with all types of drive, our algorithm was able to correctly identify the start of a chase and matched the results of visually identifying the start of the chase.

### Smoothing

In some specified figures, curves were smoothed to reduce noise from the limited number of simulations by displaying the weighted average of a data point and three adjacent points on either side (at weights of 75%, 37.5%, and 18.75%, decreasing based on distance to the center point). Near the ends in regions of rapid change, the number of data points used for smoothing on either side was equal to the number of data points between the point in question and the end point (zero for the end point) using the same weighting system.

### Data generation

Simulations were run on the computing cluster at the Department of Computational Biology at Cornell University. Data processing and analytics were performed in Python, and figures were prepared in R. All SLiM configuration files for the implementation of these suppression drives are available on GitHub (https://github.com/MesserLab/Chasing).

## RESULTS

### Dynamics of suppression drives in panmictic populations

We analyzed four different suppression drive strategies in this study. The first two strategies are homing-type drives, targeting either a female or both-sex fertility gene. The former has already been demonstrated in *Anopheles*^15^. The third is a driving Y chromosome, based on an X-shredder allele. The fourth is a TADS suppression drive^34^ (see Methods for details of the different drive mechanisms).

In our panmictic population model, each of the drives, in idealized form, quickly increased in frequency after release and eliminated the population (Figure 1A), consistent with previous findings^16,37,38^. This remained true even if we assumed somewhat imperfect drives. For example, if drive efficiency and drive fitness were reduced to 0.8, each drive still successfully eliminated the population, though somewhat more slowly. Similarly, each drive can tolerate higher low-density growth rates (values of up to 12), despite the fact that increasing this parameter makes it more difficult for a drive to ultimately eliminate a population, given that a higher growth rate allows production of larger numbers of offspring when population size is small. The only exception to this is the Driving Y, which usually fails to suppress the population when 1 / (1 − *drive efficiency*) is less than the *low-density growth rate*. This is consistent with previous findings showing that such drives reach an equilibrium population size in this case instead of completely suppressing the population, even after fixation of the drive allele^22,38,39^.

**Figure 1.**
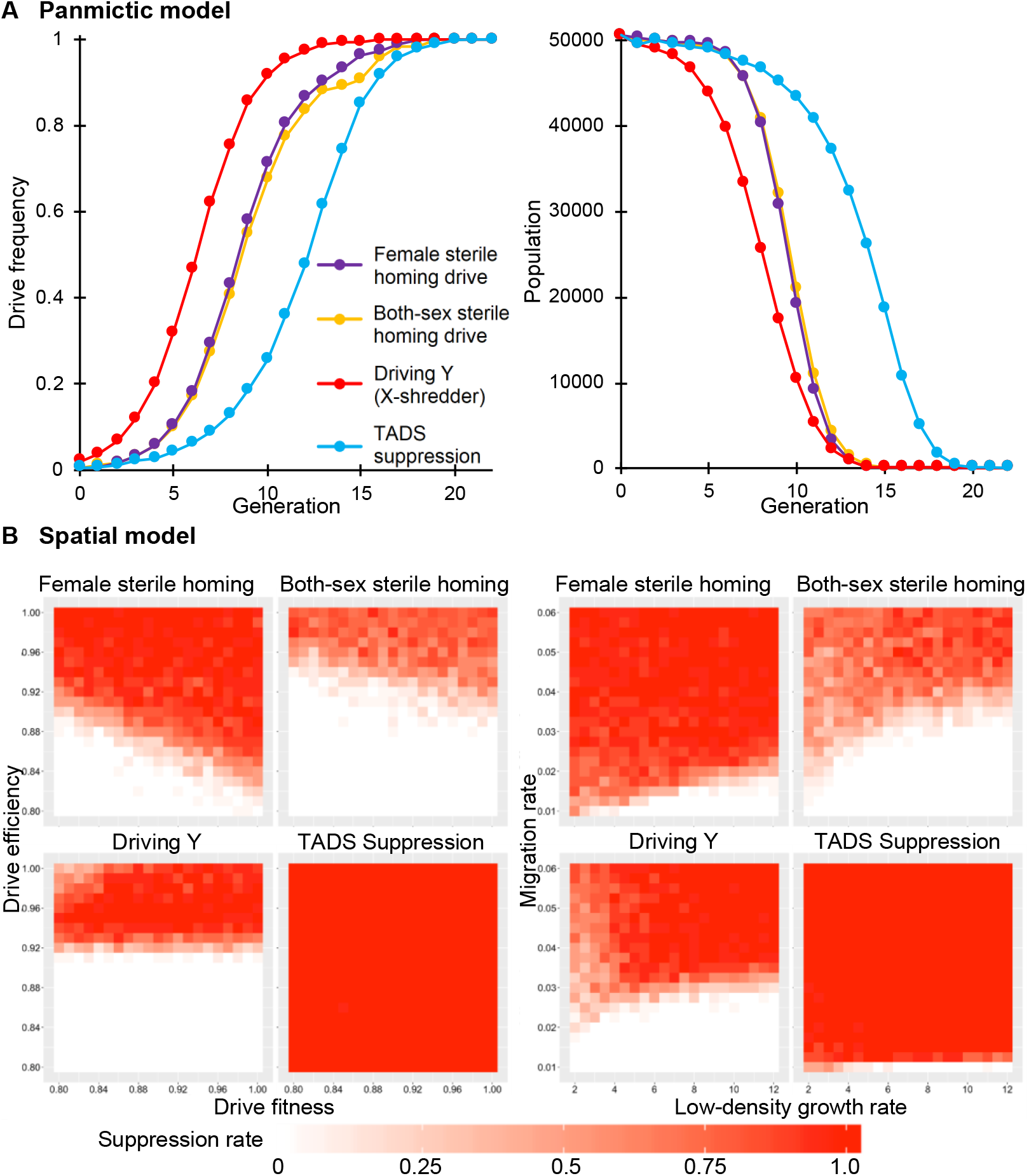
Dynamics of suppression gene drives in our panmictic and spatial models. (**A**) Drive heterozygotes (drive-carrying males for the Driving Y) were released at 1% frequency into a panmictic population of wild-type individuals, and the drive allele frequency and population size were tracked for each generation until the population size reached zero. The data displayed shows averages for 20 simulations. (**B**) Drive heterozygotes (drive-carrying males for the Driving Y) were released in a 0.01 radius circle into the middle of spatial population. Outcomes were tracked for 1000 generations for each simulation. The suppression rate specifies the proportion of simulations where the population was eliminated. Drive fitness and drive efficiency were varied on the left; low-density growth rate and migration rate were varied on the right. Each point represents the average of at least 20 simulations.

### Suppression is less effective in spatially continuous populations

Panmictic population models can help us understand the basic dynamics of a gene drive, but real-world populations are usually structured, with individuals moving over a continuous landscape. To better understand how the dynamics of a suppression drive may be affected by such factors, we implemented a spatial simulation model in which individuals inhabit a two-dimensional arena. In this model, mates are chosen locally, offspring disperse a limited distance from their parents, and population density is controlled by local competition. The level of localization can be varied in our simulations by the *migration rate* parameter, which determines both the average dispersal distance of offspring and the radius over which mates can be selected (see Methods for details).

Figure 1B shows that the ability to eliminate the population is substantially reduced in our spatial model compared to the panmictic model. As we varied drive efficiency and fitness values between 0.8 - 1.0 (representing high-efficiency drives), low-density growth rate between 2 - 12, and migration rate between 0.01 - 0.06, only the TADS drive was able to consistently eliminate the population within 1000 generations. While the female sterile homing drive generally performed better than the both-sex sterile homing drive and the Driving Y system, all three of these strategies failed over large areas of the parameter space tested. For example, none of these three drives was able to reliably eliminate the population when drive efficiency was below 0.9 or when the migration rate was below 0.03.

### Chasing dynamics accounts for majority of drive failure in the spatial model

We wanted to test what causes drive failure in the spatial model. For the Driving Y, it is known that even in panmictic models, elimination can fail when drive efficiency is low and low-density growth rate is high, despite the drive allele becoming fixed in the population^22,38,39^. This occurs when enough X chromosomes escape shredding in each generation for the resulting females to be able to maintain the population. We observed such an equilibrium in our spatial model of the Driving Y as well (Figure S1), but this mechanism does not fully explain drive failure over the whole parameter range, nor does it account for any failures of the other drive types. Another possible scenario is that failure is due to loss of the drive, which allows the wild-type population to rebound afterwards. We found that this indeed occurred in some cases (Figures S2–3), particularly for the both-sex sterile homing drive. However, this was too infrequent to account for the high failure rate.

Instead, we found that in most of the cases of drive failure, both drive and wild-type alleles coexisted in the population. Closer analysis of the spatio-temporal dynamics of drive carriers and wild-type individuals in these scenarios revealed an interesting pattern we term “chasing”, which is characterized by large fluctuations in population density over time and space (Figure 2). Chasing occurs when the drive has cleared substantial parts of the population, creating empty areas into which wild-type individuals can then escape from drive-populated areas. Because of limited competition in those areas, the wild-type population rebounds there quickly. Drive alleles then move in from the perimeter of the recolonized area, “chasing” the wild-type alleles and eventually suppressing the population in that area again. Meanwhile, wild-type alleles ahead of the drive are still recolonizing empty regions, preserving the chasing dynamics. Several videos illustrating chasing behavior in our simulations for different drive types are available on YouTube (tinyurl.com/y5vjsfy2).

**Figure 2.**
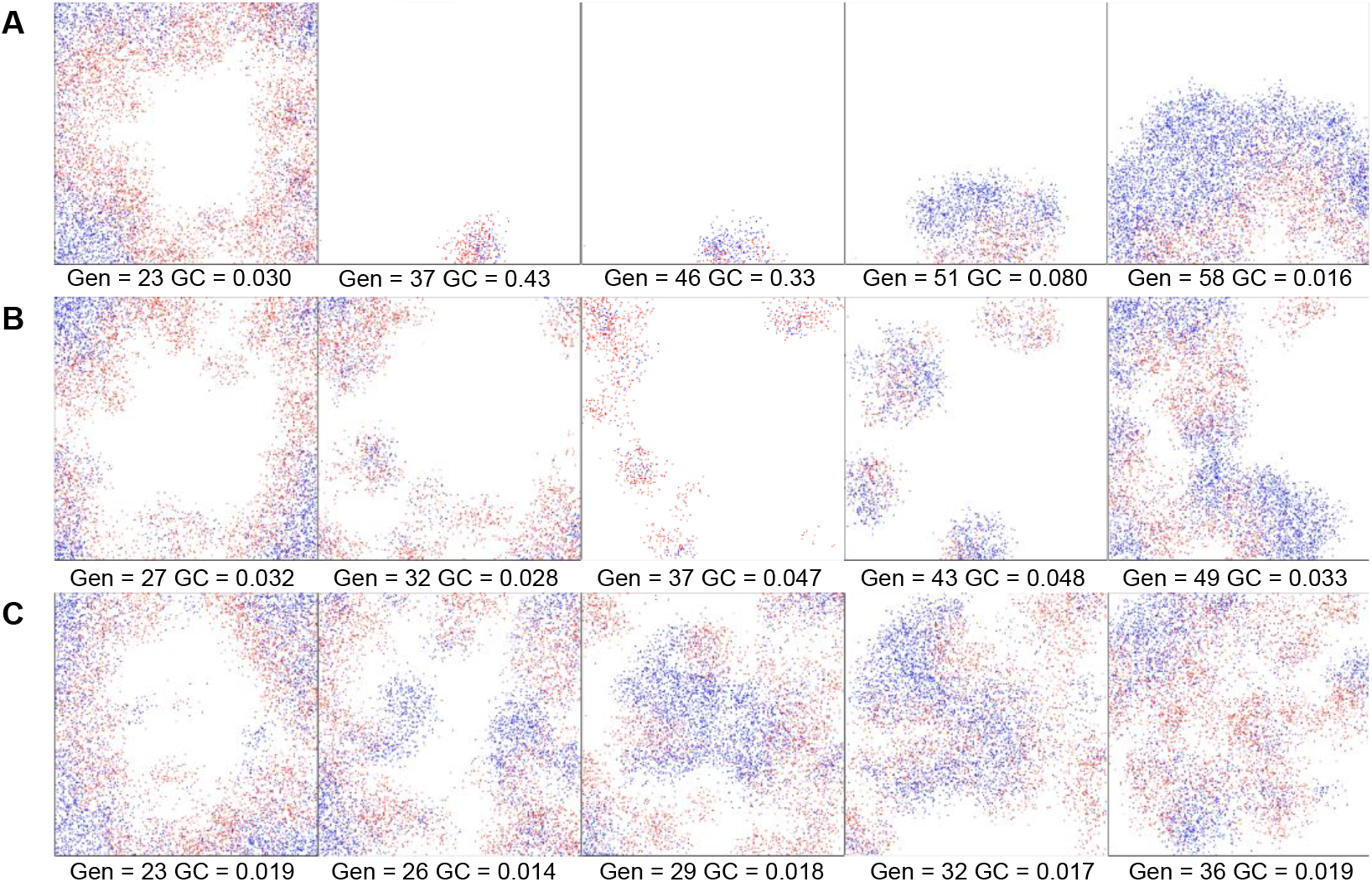
The chasing phenomenon. Snapshots of the population are shown in different generations (Gen) during a period of chasing of a female sterile homing drive. Red individuals have at least one drive allele and blue individuals have no drive alleles. Three scenarios with different parameters are shown where the chasing behavior is characterized by high (**A**), medium (**B**), and low (**C**) Green’s coefficient (GC), a measure of the degree of clustering. In (A), the drive cleared most of the area, but some wild-type individuals in a single patch persisted near the bottom of the area. These then spread into the large area of empty space, with the drive chasing them. In some cases, there can be multiple distinct chasing patches at the same time (B-C).

Importantly, chasing is different from equilibrium scenarios that can also lead to coexistence of drive and wild-type, such as observed for the Driving Y in both panmictic and spatial scenarios. In equilibrium scenarios, overall population density is uniform across space, and local drive allele frequencies are similar across all regions. By contrast, chasing is characterized by strong clustering of individuals in space, with often substantial allele frequency differences between clusters and large fluctuations in cluster sizes and locations over time and space. The availability of empty spatial areas is critical to bring about these dynamics. Thus, chasing is fundamentally a spatial phenomenon that cannot occur in panmictic models.

To detect whether chasing has occurred at any point in a given simulation run, we developed a statistical test based on the longitudinal analysis of population size changes and measures of spatial clustering (see Methods). We find that chasing is generally common in areas of the parameter space where the drive struggles to eliminate the population (Figures S4–5), though these ranges do not overlap exactly (compare with Figure 1B).

### Parameters affecting drive outcomes in the spatial model

To better understand the factors that determine the probability of different drive outcomes in light of the chasing phenomenon, we analyzed each drive type, varying one parameter at a time. The outcomes of simulation runs were divided into six possible categories: (i) population elimination without prior chasing; (ii) elimination after chasing; (iii) establishment of a stable equilibrium with reduced population size but where chasing did not occur (applicable only to the Driving Y); (iv) long-term chasing (simulations in which chasing was still occurring after 1000 generations); (v) loss of the drive after a chase; (vi) loss of the drive without prior chasing.

Figure 3 shows the results of these analyses, revealing complex dependencies of drive outcomes on individual parameters and pronounced differences between drive types. In general, the TADS suppression drive was the only system that remained effective across the full range of parameters tested, except when migration rate was very low. The female-sterile homing drive also performed well but was still unable to induce population elimination for much of the parameter space. The Driving Y and both-sex sterile homing drives were generally less effective. The both-sex sterile homing drive in particular showed low performance since it was often lost from the population, usually after an initial period of chasing. A variant of this drive that induces lethality instead of sterility showed broadly similar performance (Figure S6, Supplementary Results).

**Figure 3.**
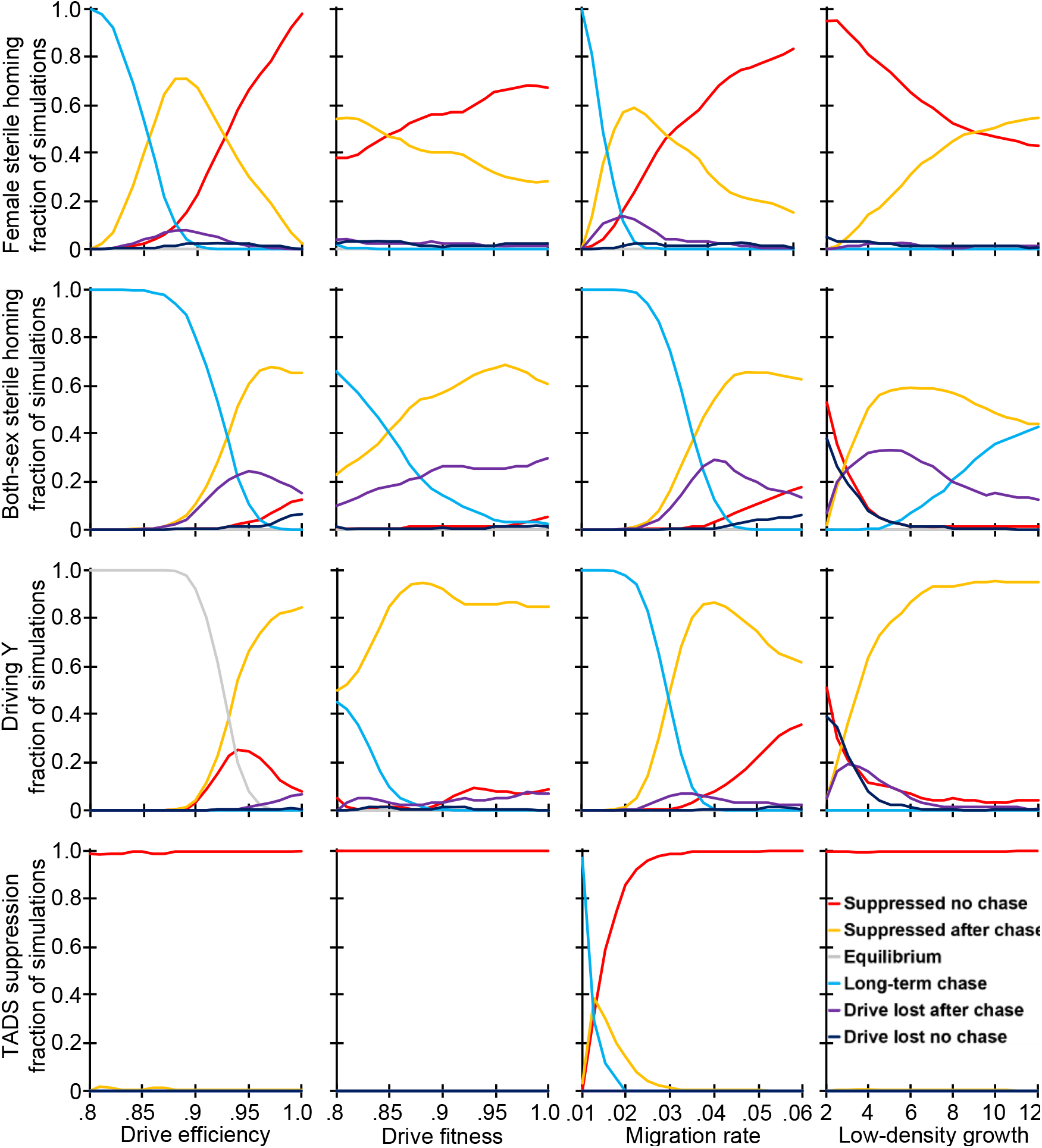
Effects of drive and ecological parameters on suppression outcomes in the spatial model. Drive heterozygotes (drive-carrying males for the Driving Y) were released into the middle of a wild-type population. The proportion of different simulation outcomes is shown (“long-term chase” represents continued chasing behavior at generation 1000). Curves were obtained by averaging at least 100 simulation runs for each tested parameter value and then smoothed as described in the methods to reduce noise.

Drive efficiency had a dramatic effect on the success rate, even though we only considered drives with efficiency levels at or above 80% in the first place. For the female-sterile homing drive, as efficiency increased, long-term chasing outcomes were replaced with elimination after chasing, which in turn were eventually replaced with elimination without chasing. For the both-sex sterile drive, as efficiency increased, long-term chasing outcomes became less common and elimination after chasing became more common. However, the rate at which the drive was lost also increased before declining again when drive efficiency approached 100%. For the Driving Y, higher drive efficiency prevented equilibrium outcomes (when the drive fixated, but did not chase or eliminate the population), but even for its optimal efficiency, which was somewhat below 100%, this drive did not achieve elimination in all simulations, unlike TADS or the female sterile homing drive. In some simulations at this optimal level, elimination occurred quickly, but usually it was after a period of chasing. Increasing drive fitness generally shifted outcomes toward higher elimination rates for all drives, but this effect was of considerably lower magnitude than the effect of increasing drive efficiency in the parameter space we considered.

In addition to drive parameters, ecological parameters also had notable effects on outcome rates. Increasing the migration rate generally increased the rate of elimination outcomes, consistent with the fact that higher migration should make the spatial model more similar to a panmictic model. Higher low-density growth rates generally decreased the rate of elimination outcomes while increasing the rate of chasing outcomes, though the rate at which the drive was lost was also reduced. The overall population density had little effect on outcomes (Figures S7, Supplementary Results). However, we found that changing the boundary of our arena to an unbounded toroidal space decreased the rate of successful elimination after chasing (Figure S8, Supplementary Results).

### Impact of chasing on suppression potential

To better understand the impact of chasing on the overall goal of population suppression, we first examined the duration of chasing in runs where the population was eliminated after a chase (Figures S9–10). We found that in areas of the parameter space where chasing was common, the time interval of chasing tended to be longer, usually several hundred generations. Where chasing was less common, the time interval of chasing tended to be shorter, comprising only a few generations before complete suppression.

The objective of a suppression drive could still be partially fulfilled even without achieving elimination if the population size is sufficiently reduced. This often occurs in chasing scenarios. To determine the magnitude of population reduction, we analyzed the average population size during chasing, regardless of final outcome, over a range of parameters (Figures S11–12). In general, the migration rate had a dominant effect, and when chasing was more common, the average population size during chasing was typically higher. Nonetheless, population reductions by a factor of 2-3 were common. When chasing was less common, the average population size was even smaller. The average Green’s coefficient during the chase was found to be lower when chasing was more common, indicating the presence of a greater number of chasing clusters at any given time (Figures S13–14, Supplementary Results).

### The effect of inbreeding on chasing

Previous studies have found that inbreeding can pose a substantial obstacle to the spread of a gene drive^26,40^. To test whether these results extend to our spatial model and explore possible connections to the chasing phenomenon, we studied how varying the probability of matings between siblings affected drive outcomes in our spatial model. (Figure 4). Consistent with previous results, we found that increased inbreeding (achieved in our model by increasing the preference for choosing a sibling as a mate) resulted in a reduced likelihood of elimination and more chasing. Similarly, if individuals had a reduced sibling mating rate, successful elimination became more likely. This was also observed when we reduced the fecundity of matings between siblings, representing the effects of inbreeding depression (Figures S15–16).

**Figure 4.**
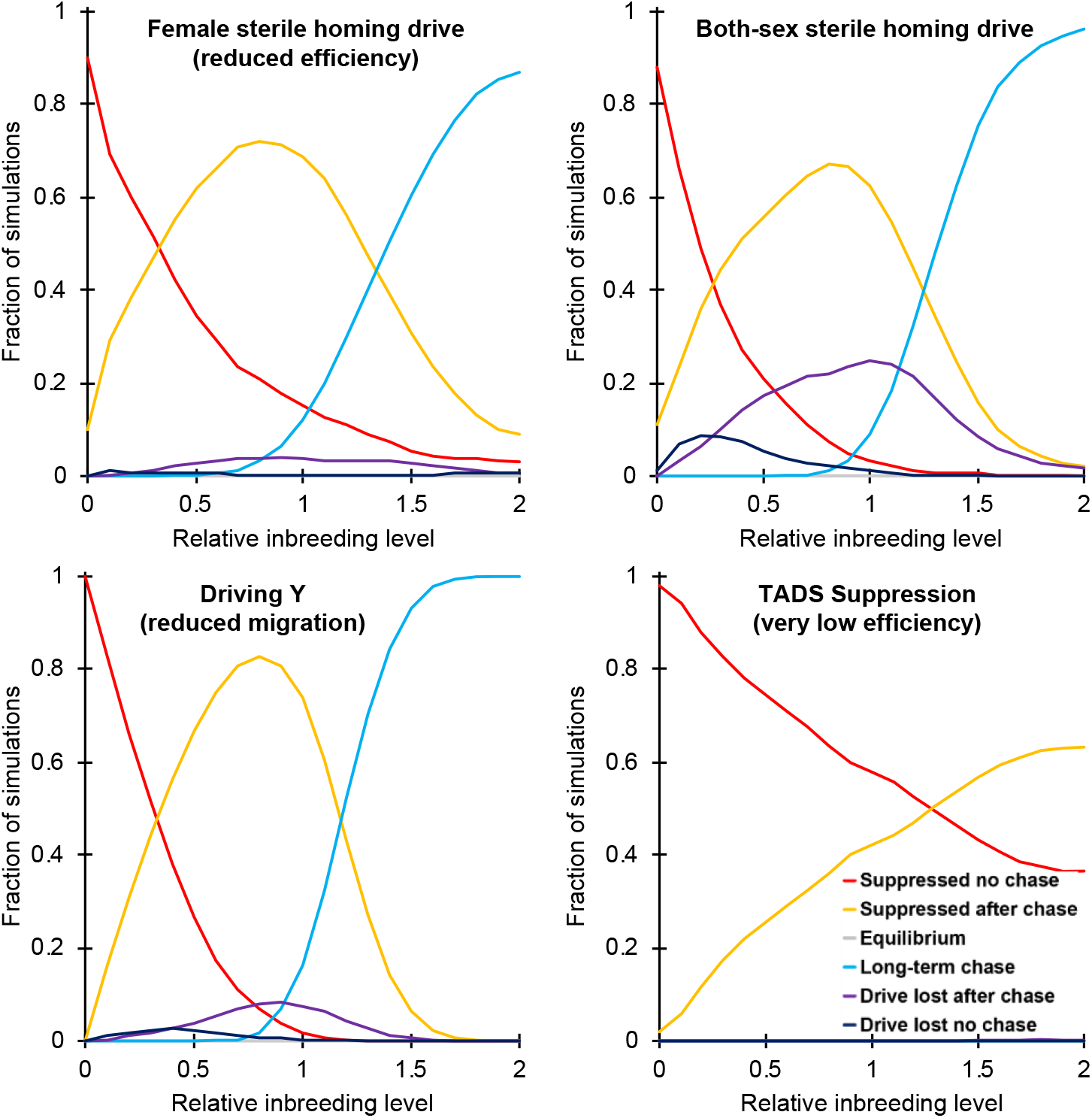
The effect of inbreeding on suppression outcomes in continuous space. Drive heterozygotes (drive-carrying males for the Driving Y) were released into the middle of a wild-type population. The proportion of different simulation outcomes is shown. The relative inbreeding level specifies the preference a female gives to siblings when choosing a mate as compared to non-siblings (a value of 1 means that no preference is given, while a value of 2 means that siblings are twice as likely to be chosen), before adjustment by fitness. To show a greater dynamic range of outcomes, some default parameters were modified (female sterile homing drive: efficiency and fitness was reduced to 0.92, migration rate was reduced to 0.035, and low-density growth rate was increased to 8; driving Y: migration rate was reduced to 0.0325; TADS suppression drive: efficiency and fitness was reduced to 0.8, migration rate was reduced to 0.02, and low-density growth rate was increased to 12). Curves were obtained by averaging at least 100 simulation runs for each tested parameter value and then smoothed as described in the methods to reduce noise.

### Chasing can lead to drive failure by resistance allele formation

Cleavage repair by end-joining or incomplete homology-directed repair can lead to the formation of resistance alleles that do not match the drive’s gRNAs and are thus immune to future cleavage^41–43^. Thus far, we only considered resistance alleles that disrupt the target gene function, which usually do not have a drastic impact on the success rate of a suppression drive. However, some resistance alleles could preserve the function of the target gene and ultimately stop a suppression drive from spreading. Such function-preserving mutations are known as “r1” resistance alleles. A recent study in *Anopheles* was able to prevent the formation of r1 alleles in small population cages^15^, but it is unclear exactly how well the formation of such alleles can be mitigated.

To test the potential impact of r1 resistance alleles on drive outcomes, we varied the r1 resistance rate for the two drives in our model that could be prone to such alleles (the both-sex and female sterile homing drives) (Figure 5A). The r1 rate here specifies the fraction of resistance alleles that become r1 alleles. We did not see an impact of resistance in our model with *K* = 50000 individuals for r1 rates below 10^−5^, while for rates above 10^−3^, outcomes were dominated by resistance. Notably, in drive failures at intermediate r1 rates, the first r1 alleles usually arose well after the drive had started chasing, suggesting that if chasing had not occurred, such alleles would likely not have been able to prevent elimination.

**Figure 5.**
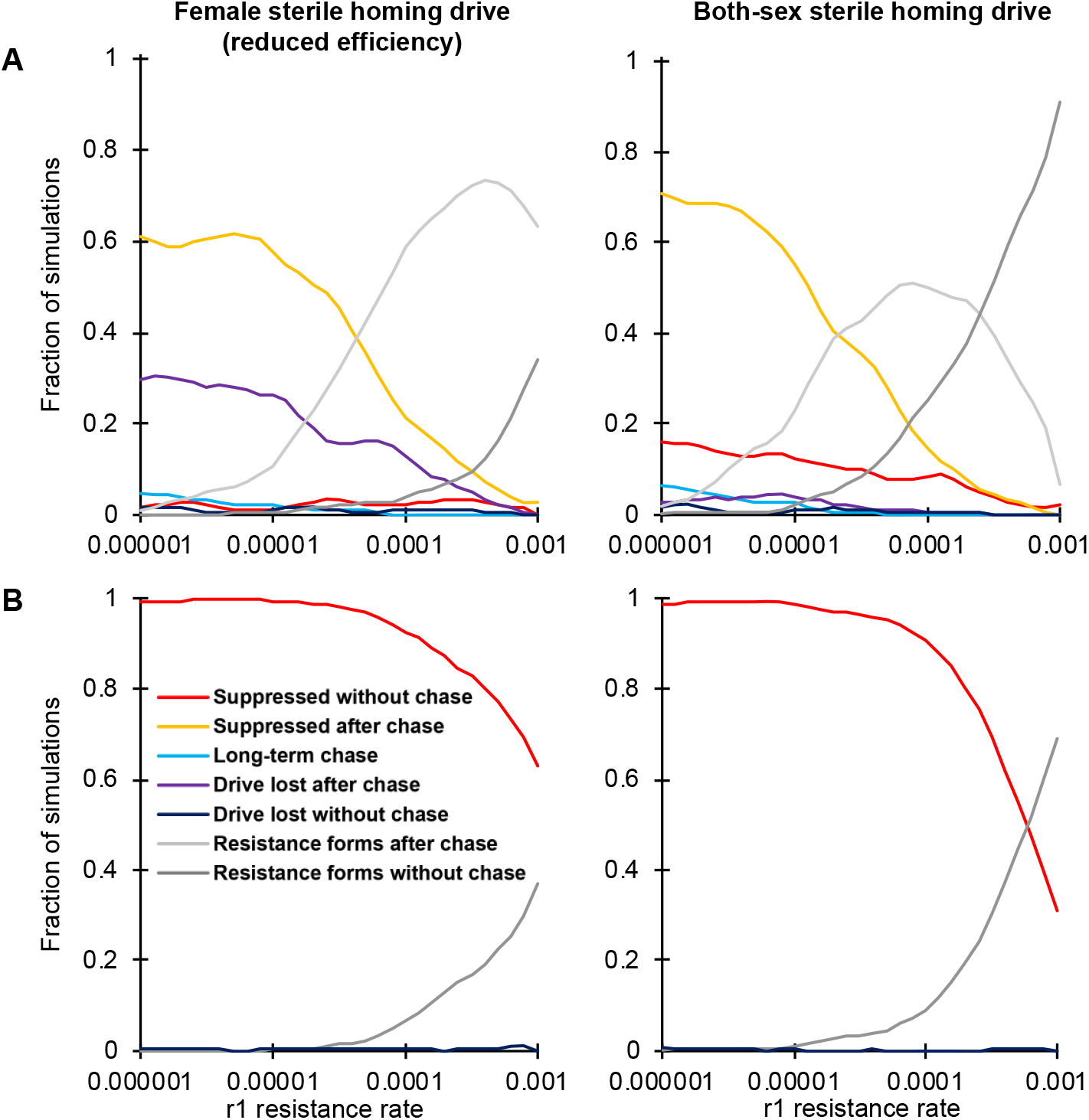
The effect of resistance allele formation on suppression outcomes. Drive heterozygotes (drive-carrying males for the Driving Y) were released into a (**A**) spatial or (**B**) panmictic population. The proportion of different outcomes is shown for the female sterile homing drive and the both-sex sterile homing drive. Resistance outcomes refer to simulations where resistance alleles that preserve the function of the target gene reached at least 10% frequency with at least 500 individuals present. Each point represents the average of at least 100 simulations. The r1 rate is the fraction of resistance alleles that preserve the function of the target gene. To better show a range of outcomes, some default parameters were modified for the female sterile homing drive: efficiency and fitness was reduced to 0.92, migration rate was reduced to 0.035, and low-density growth rate was increased to 8. Smoothing of curves was performed as described in the methods to reduce random noise.

This underscores the importance of chasing dynamics. Even temporary chasing raises the effective number of wild-type alleles that must be converted by the drive before successful elimination could occur, with each conversion possibly resulting in an r1 allele that may ultimately prevent elimination and allow the population to rebound. This is in stark contrast to panmictic models of these drives (Figure 5B), where r1 alleles can only thwart drive systems when occurring at much higher rates.

## DISCUSSION

In this study, we demonstrated that suppression gene drives can frequently experience chasing dynamics in spatial population models, which often prevents complete population elimination. This phenomenon occurs when wild-type individuals move into areas previously cleared by the drive, where they can rebound quickly due to reduced competition. Drive alleles then follow and chase wild-type alleles across the landscape.

The consequences of chasing could potentially impact the decision to deploy a suppression drive. At minimum, chasing may delay complete suppression, often by a substantial time interval. More robust chasing can persist perpetually, depending on drive and environmental parameters. Furthermore, the larger sizes of most realistic populations compared to those we modeled provide more possibilities for a chasing situation to start. The overall population size can still be reduced substantially during chasing, yet continuous conversion of wild-type alleles may eventually result in the formation of functional resistance alleles. If such alleles do arise, population size can rebound quickly.

The definition of chasing we employed in this study is qualitative in nature, and we hope that future studies can develop a formal definition that would allow us to better understand how and when chasing is initiated. Simulations in one-dimensional space suggest that a chase can start by local elimination of the drive, thereby opening a migration route for wild-type individuals to recolonize empty areas. However, it is also possible for wild-type individuals to directly permeate an expanding wave of the drive to reach empty areas behind the wave that have been cleared by the drive (Figures S17–22, Supplementary Results).

Chasing dynamics for a spatial suppression drive are similar to other phenomena that have previously been described. The dynamics between areas with drive individuals, wild-type individuals, and empty space are similar to the elements in the game of “rock-paper-scissors”, in which no one element always dominates. Such dynamics have been observed in other biological systems such as coral reef invertebrates^28^ and bacterial populations^29^. In the latter case, three cyclically-competing strains were able to form coexisting patches, with each patch chasing another type. However, when the population was well-mixed (analogous to our panmictic model), one strain always dominated^29^. In models of predator-prey interactions, continuous space was also essential for predators, prey, and empty space to coexist^30^.

Outcomes consistent with chasing behavior have already been observed in other models of suppression gene drives in structured populations^22,24,26^. In these studies, it was generally suggested that the drive efficiency should be high enough to avoid drive loss due to drift but should not be so high that the drive eliminates patches of wild-type and itself before being able to spread to an adjacent region. This contrasts with our findings in continuous space, where except for the Driving Y, drives with maximum efficiency generally were most effective.

Compared to our continuous-space model, the amount of chasing could possibly be increased by long-distance dispersal (by water, wind, human transportation, etc.), as this could make it easier for wild-type individuals to escape into empty areas far away from any drive-carrying individuals, where they could then expand quickly. Environmental variation may also facilitate chasing if regions exist with ecological parameters that facilitate chasing, which could then serve as “seeds” for recurrent, temporary expansions of wild-type individuals into surrounding areas that are less amenable to chasing.

We found that the propensity of chasing is substantially affected by both drive performance and ecological parameters. In general, even modest reductions in fitness and efficiency greatly increased the likelihood of chasing and reduced the chance of successful elimination. However, the optimal efficiency for the Driving Y (X-shredder) was somewhat less than 100%, as seen in a previous study^24^. In our model, this may be because the drive suppresses rapidly, and slightly reduced efficiency could prevent local stochastic loss of the drive, thereby making it harder for wild-type individuals to pass the drive and recolonize previously cleared empty areas to start a new chase. In simulations where the low-density growth rate was very low, we often saw a stochastic loss of the drive when the numbers of drive and wild-type individuals had reached low levels. Otherwise, this parameter did not have a large effect. Higher migration rates shifted the range of outcomes in favor of elimination and reduced chasing propensity, consistent with the fact that this should generally shift the spatial model closer to the panmictic model.

The drive types we investigated had markedly different effectiveness in their ability to suppress populations in continuous space. Understanding the underlying reasons for this poses an interesting topic for future study. Our initial analysis suggests that stochastic factors and the “thickness” of the advancing drive wave may play a key role in these differences (Table S1, Figures S23–25, Supplementary Results). One clear conclusion is that homing-type suppression drives should be targeted to a recessive gene that affects only one sex (such as a female fertility gene), given that the both-sex drive had a substantially higher tendency to chase and also suffered from higher stochastic loss of the drive in our spatial model. The female fertility drive, on the other hand, performed quite well when it had high efficiency, which is promising given that such drives have already been constructed in *Anopheles gambiae*^15^. The X-shredder (a type of driving Y chromosome and very similar to a TADS-based driving Y^34^) also performed worse than the female sterile homing drive. TADS suppression had the highest effectiveness of all drive types we tested. If suitable gene targets for such a system can be identified, this could enable the development of drives that can both minimize resistance alleles with multiplexed gRNAs (a useful strategy but with substantial limitations in homing-type drives^31^) and achieve effective suppression over a large range of parameters.

Evolution of an increased tendency for inbreeding has been suggested as a mechanism by which populations could avoid the suppressive effects of a gene drive^26^, and our studies in continuous space support this notion. We found that higher levels of inbreeding can indeed substantially reduce the effectiveness of the drive by increasing the likelihood of chasing. However, inbreeding avoidance (or inbreeding depression) can actually work in favor of the drive. In many real-world target populations, these latter effects could play an important role in determining the likelihood of drive success, and since suppression could occur rapidly, there may be insufficient time for the evolution of strategies that increase inbreeding in the population.

Overall, we have shown that suppression gene drives can exhibit rich dynamics in spatially continuous populations with a wide range of possible outcomes. In particular, the chasing effect could be a primary means by which a population can escape elimination by a drive. Thus, to accurately predict the outcome of a suppression strategy, detailed models should be utilized that incorporate spatial structure and realistic migration patterns of individuals over both small and large scales, as well a variety of other ecological and demographic parameters.

## ACKNOWLEDGEMENTS

Thanks to Sandra Lapinska for assistance in generating SLiM configuration scripts and analyses. This study was supported by funding from New Zealand’s Predator Free 2050 program under award SS/05/01 to PWM and National Institutes of Health awards R21AI130635 to JC, AGC, and PWM, award F32AI138476 to JC, and award R01GM127418 to PWM.

## SUPPLEMENTARY RESULTS

**Figure S1.**
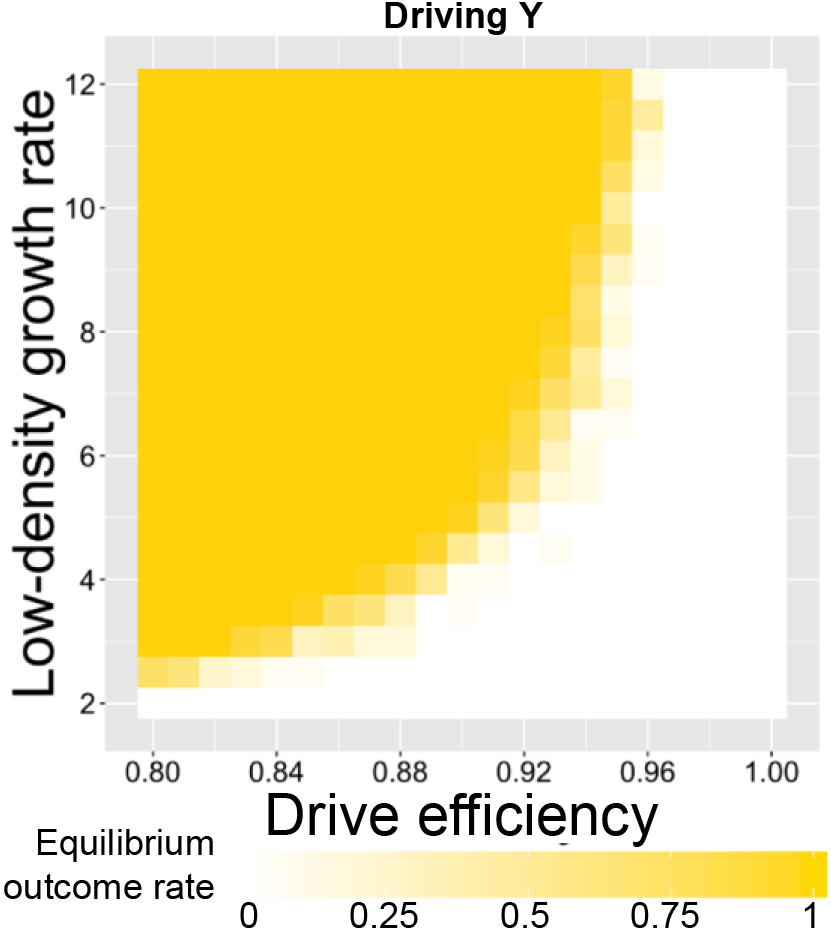
The equilibrium rate for the Driving Y in our spatial model. Drive-carrying males were released into the middle of a wild-type population in a 1×1 square. The equilibrium rate specifies the proportion of simulations where the population reached a stable equilibrium in which individuals are spread across the entire population but at a lower density than the initial wild-type population. Each point represents the average of at least 20 simulations.

**Figure S2.**
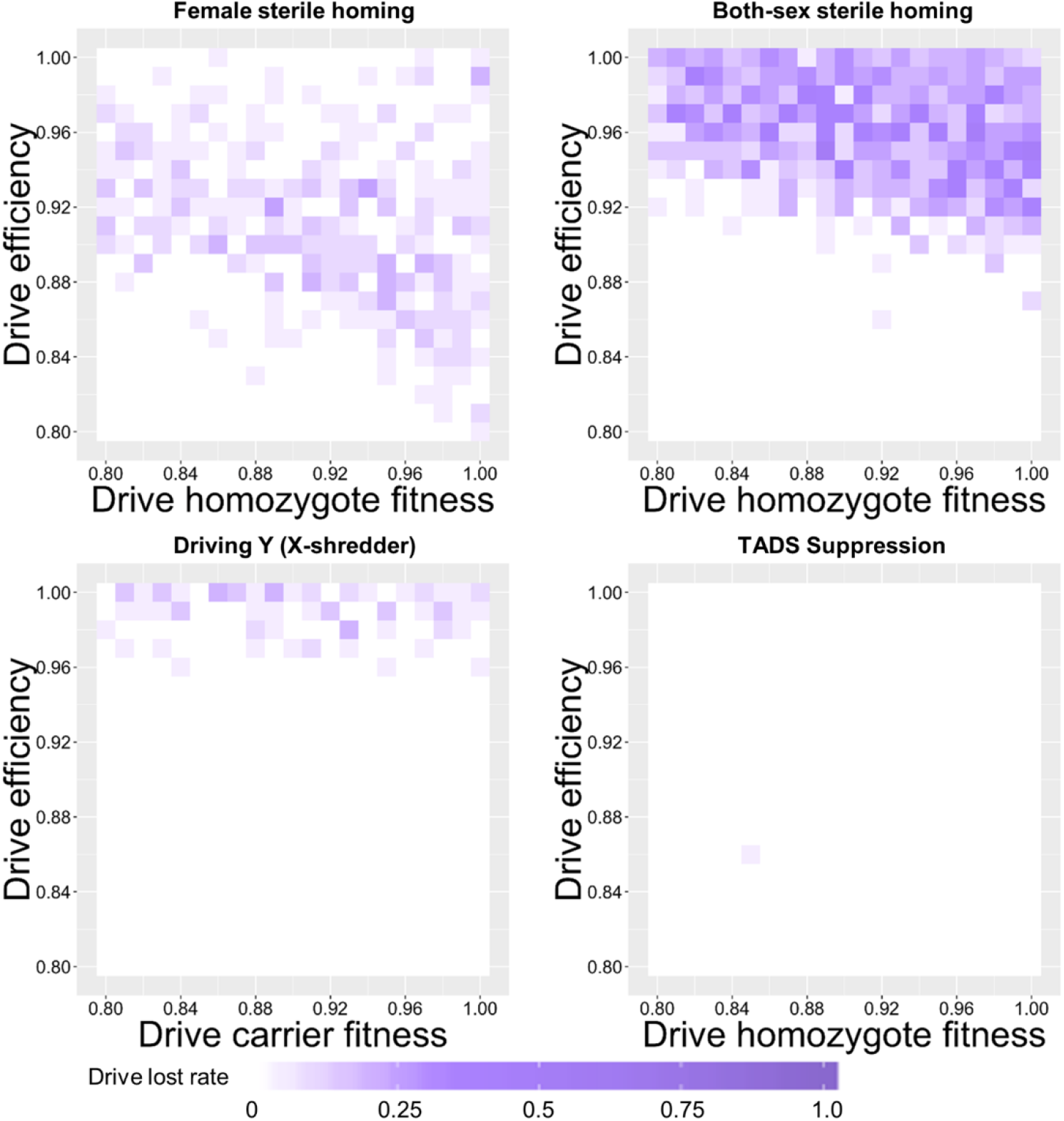
The effect of drive fitness and efficiency on the rate of drive loss in our spatial model. Drive-carrying individuals were released into the middle of a wild-type population in a 1×1 square. The drive loss rate specifies the proportion of simulations where the drive was lost. Each point represents the average of at least 20 simulations.

**Figure S3.**
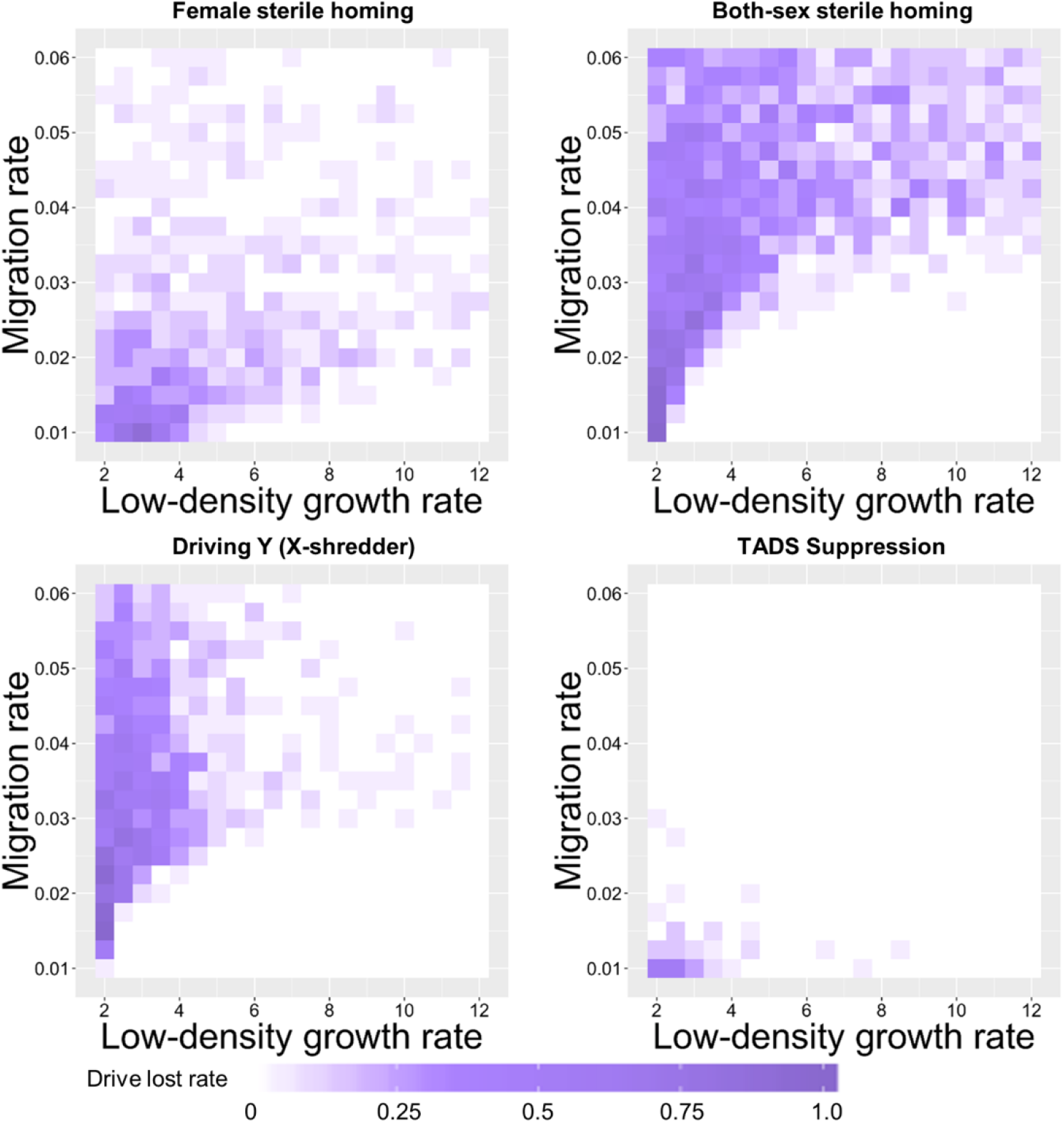
The effect of the low-density growth rate and migration rate on the rate of drive loss in our spatial model. Drive-carrying individuals were released into the middle of a wild-type population in a 1×1 square. The drive loss rate specifies the proportion of simulations where the drive was lost. Each point represents the average of at least 20 simulations.

**Figure S4.**
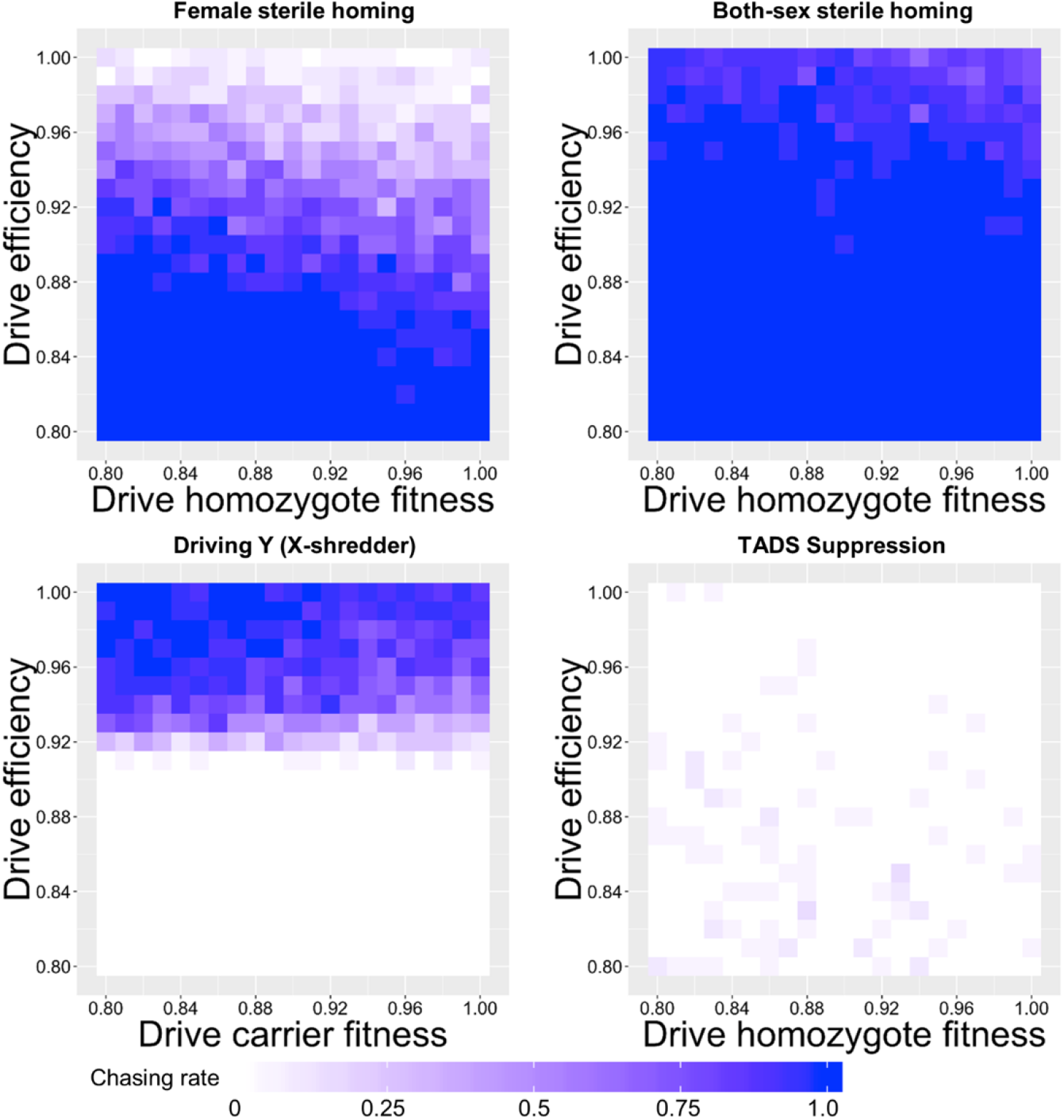
The effect of drive fitness and efficiency on the rate of chasing in our spatial model. Drive-carrying individuals were released into the middle of a wild-type population in a 1×1 square. The proportion of simulations where a chase occurred is shown. Each point represents the average of at least 20 simulations.

**Figure S5.**
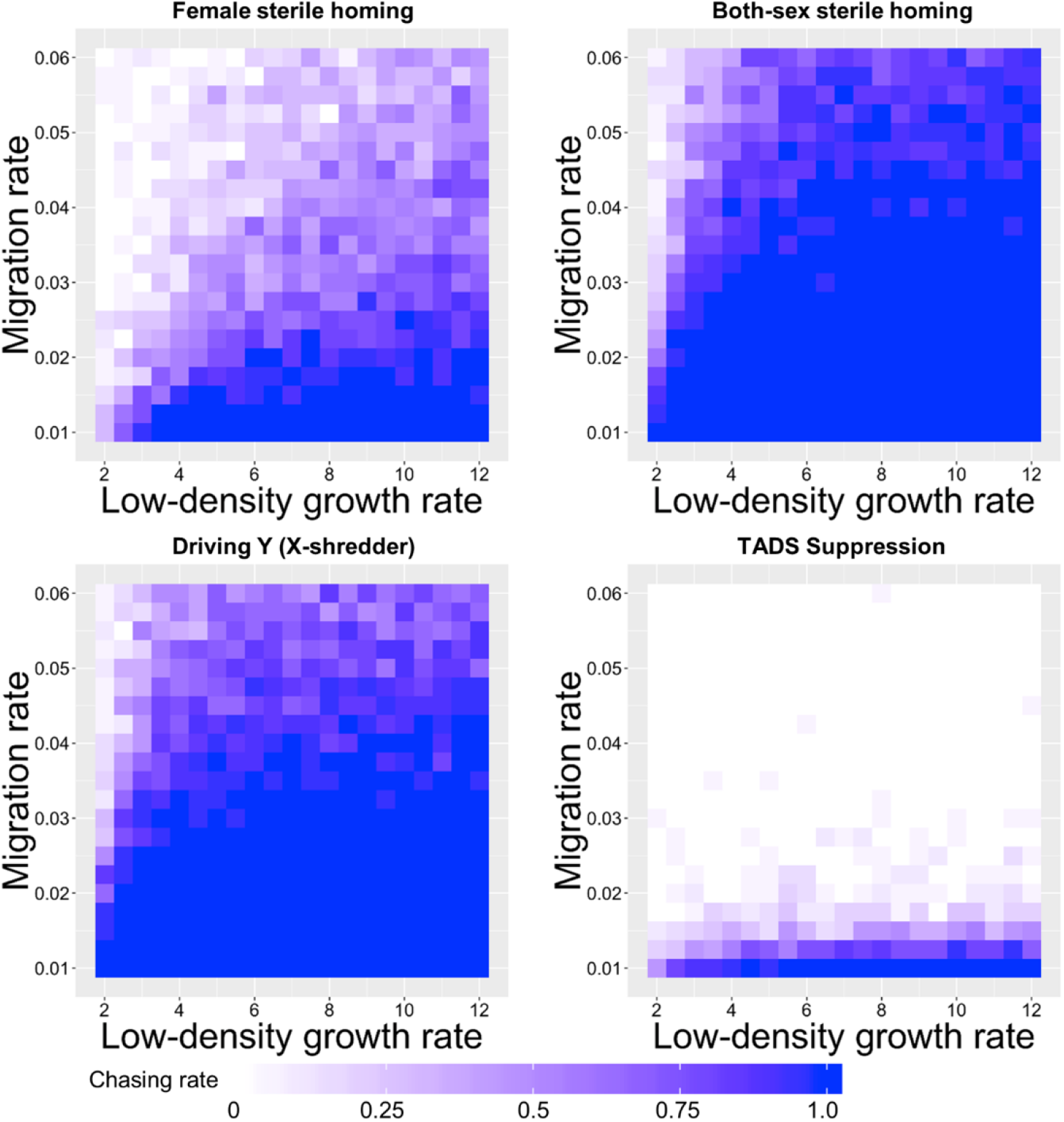
The effect of the low-density growth rate and the migration rate on the rate of chasing in our spatial model. Drive-carrying individuals were released into the middle of a wild-type population in a 1×1 square. The proportion of simulations where a chase occurred is shown. Each point represents the average of at least 20 simulations.

### Both-sex sterile vs. both-sex lethal homing drive performance

Another possible type of drive that could likely be constructed without great difficulty is a both-sex lethal homing drive. Indeed, the haplosufficient but recessive lethal targets needed for such a drive would likely be easier to find than targets for female or both-sex sterility. Comparing such a drive to the both-sex sterile homing drive, we find broadly similar performance with a few notable differences (Figure S6). Because all surviving individuals are potentially capable of producing offspring in the both-sex lethal drive (compared to the sterile drive where many matings will involve a sterile individual), stochastic effects are somewhat reduced. This likely contributes to the lower rates of drive loss compared to the both-sex sterile homing drive when the low-density growth rate is very low and proportionally higher rates of both chasing and successful suppression. However, for higher low-density growth rates, as long as migration is also at a moderate or higher level, the both-sex sterile drive has a higher rate of successful suppression, possibly due to higher suppressive power from sterile individuals mating with wild-type individuals, reducing the number of wild-type offspring. The both-sex lethal drive suppresses the population more often than the both-sex sterile drive when drive efficiency is nearly perfect. However, at a somewhat lower level of drive efficiency, the both-sex sterile drive has a greater success rate.

**Figure S6.**
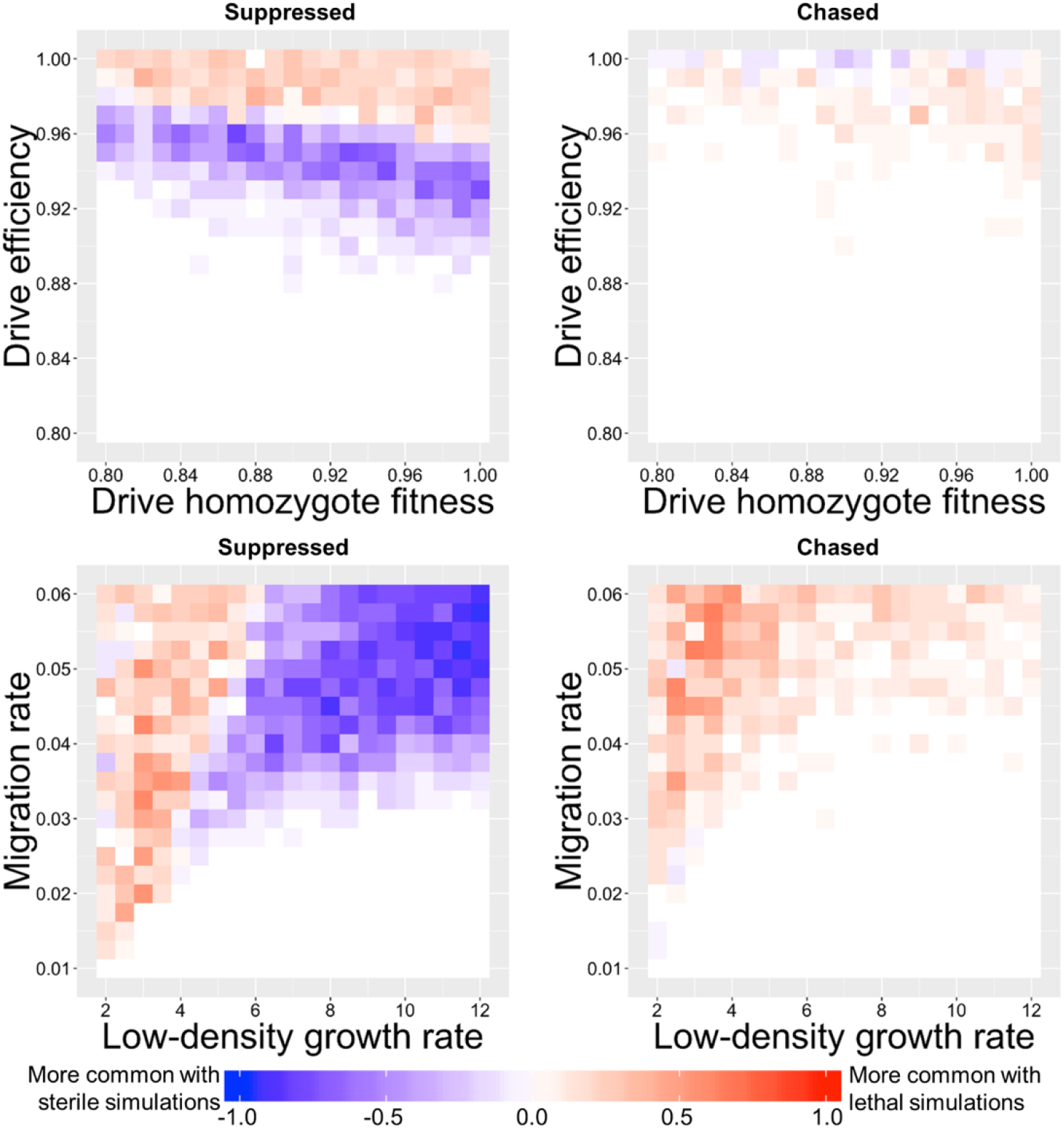
The effect of using a both-sex sterile drive versus a both-sex lethal drive on suppression outcomes. Drive-carrying individuals were released into the middle of a wild-type population in a 1×1 square. The difference in the proportion of outcomes between the both-sex sterile or the both-sex lethal drive simulations is shown. Each point represents the average of at least 20 simulations for each drive.

### Population boundaries inhibit chasing

Our square arena may be more conducive to successful suppression due to its small size and the resulting possibility that drive individuals can “corner” wild-type individuals at the boundaries of the arena, preventing easy escape to empty areas. To test this, we conducted simulations in an unbounded toroidal space and compared the results to our bounded arena using the female sterile homing drive (Figure S7). We found that chasing was initiated at about the same rate in both situations, but that successful elimination was indeed somewhat less likely to occur in the toroidal space.

**Figure S7.**
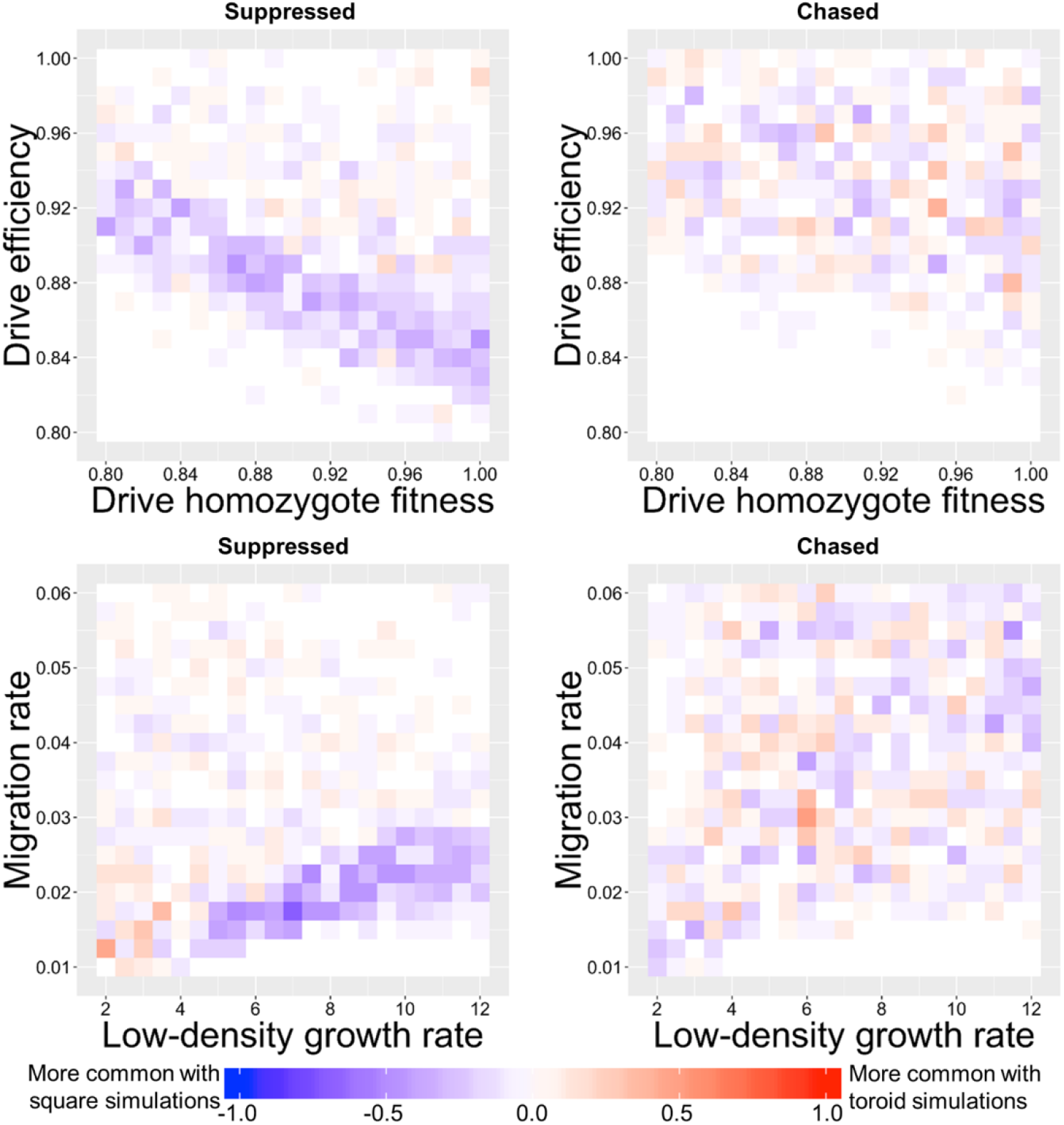
The effect of arena boundaries on suppression outcomes. Drive-carrying individuals were released into the middle of a wild-type population in a 1×1 square or a 1×1 torus. The difference in the proportion of outcomes between the square and toroidal simulations is shown. Each point represents the average of at least 20 simulations for each type of space.

### Effect of population density on suppression drive outcomes

We analyzed the influence of overall population density on drive outcomes in our spatial model but found little to no effect. This is in contrast to a previous study in which density had a strong effect on the outcome, with elimination becoming less likely at low density^22^. This may be because connectivity between regions was reduced in this study at lower density, isolating a drive to a particular region^22^. To further test for a potentially weaker effect of population density on drive outcome in our model, we adjusted the parameters of several of the drives in an attempt to make their outcomes more sensitive to such variation (Figure S8). We still found little change for the homing drives, but the modified versions of the Driving Y and the TADS suppression drive did have notably higher rates of chasing at low densities.

**Figure S8.**
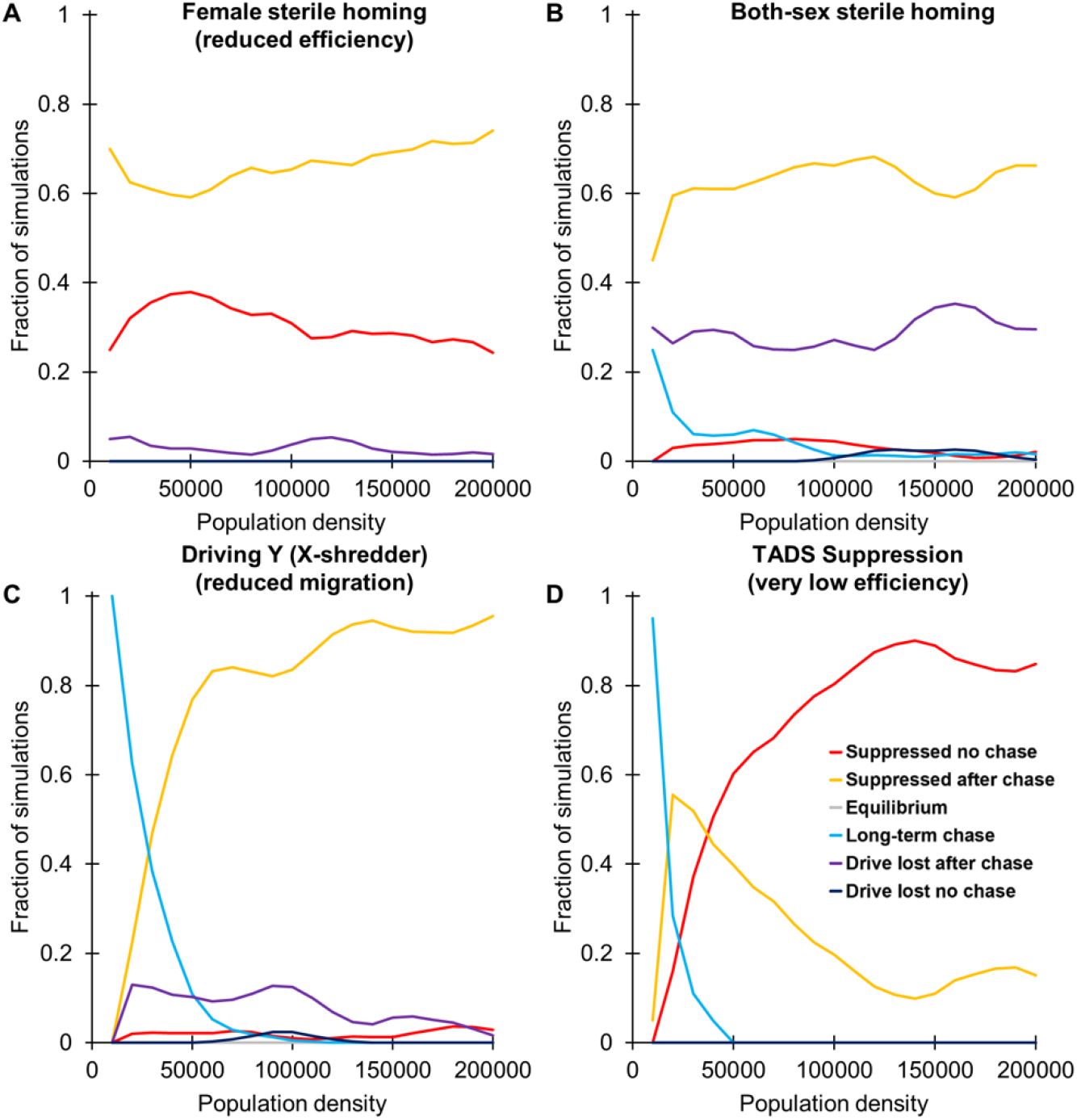
The effect of population density on suppression outcomes in the spatial model. Drive heterozygotes (drive-carrying males for the Driving Y) were released into the middle of a wild-type population. The proportion of different simulation outcomes is shown. Curves were obtained by averaging at least 100 simulation runs for each tested parameter value and then smoothed as described in the methods to reduce noise. To show a greater dynamic range of outcomes, some default parameters were modified (female sterile homing drive efficiency and fitness was reduced to 0.92, migration rate was reduced to 0.035, and low-density growth rate was increased to 8; the driving Y migration rate was reduced to 0.0325; TADS suppression drive efficiency and fitness were reduced to 0.8, migration rate was reduced to 0.02, and the low-density growth rate was increased to 12).

**Figure S9.**
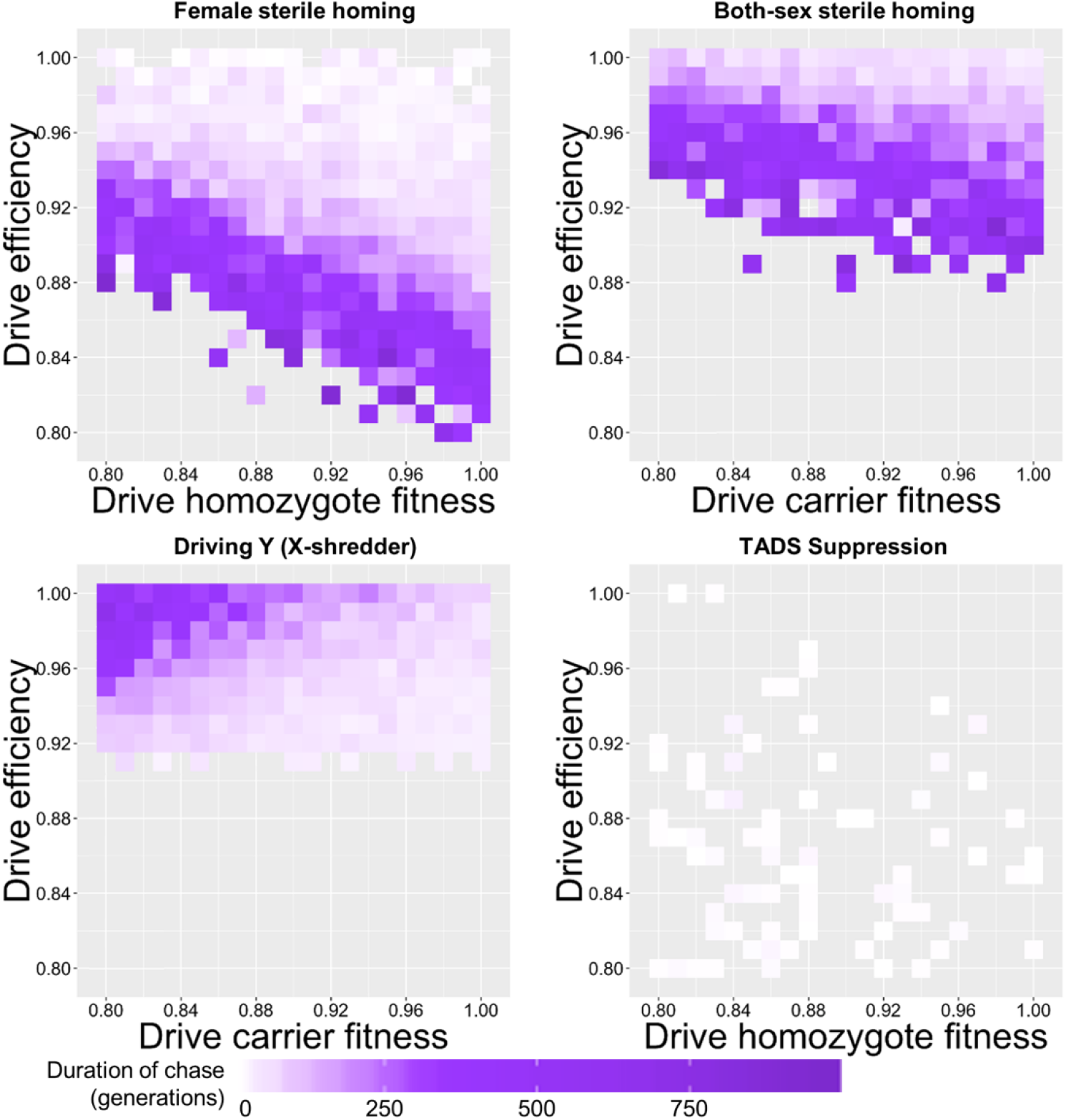
The effect of drive fitness and efficiency on the duration of chasing in continuous space. Drive-carrying individuals were released into the middle of a wild-type population in a 1×1 square. The average number of generations during which a chase occurred before suppression or loss of the drive is shown. Each point represents the average of at least 20 simulations, but only simulations that contained a chase that ended were included. Grey areas represent parameter sets in which chasing did not occur in any simulation or in which chasing occurred but did not end.

**Figure S10.**
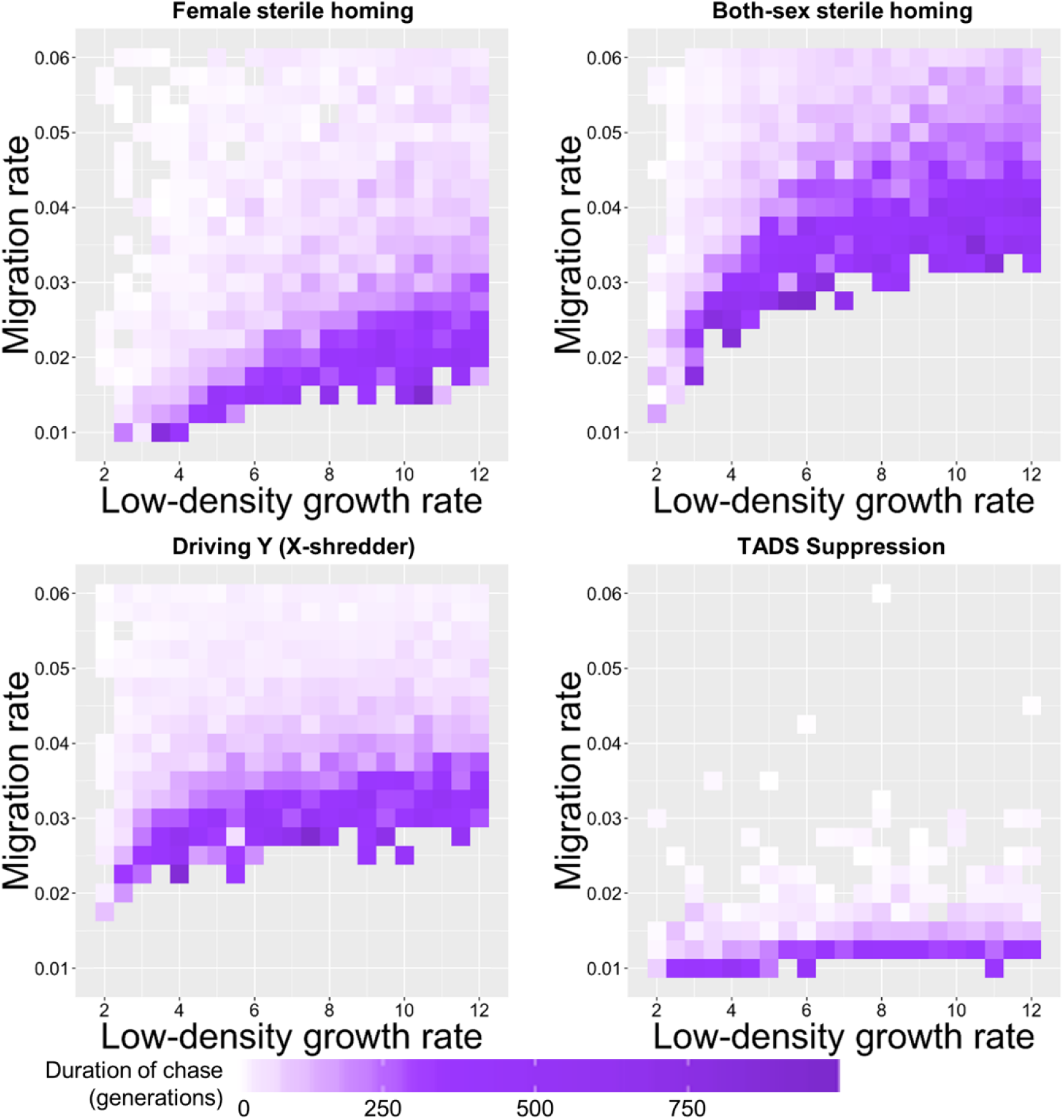
The effect of the low-density growth rate and the migration rate on the duration of chasing in continuous space. Drive-carrying individuals were released into the middle of a wild-type population in a 1×1 square. The average number of generations during which a chase occurred before suppression or loss of the drive is shown. Each point represents the average of at least 20 simulations, but only simulations that contained a chase that ended were included. Grey areas represent parameter sets in which chasing did not occur in any simulation or in which chasing occurred but did not end.

**Figure S11.**
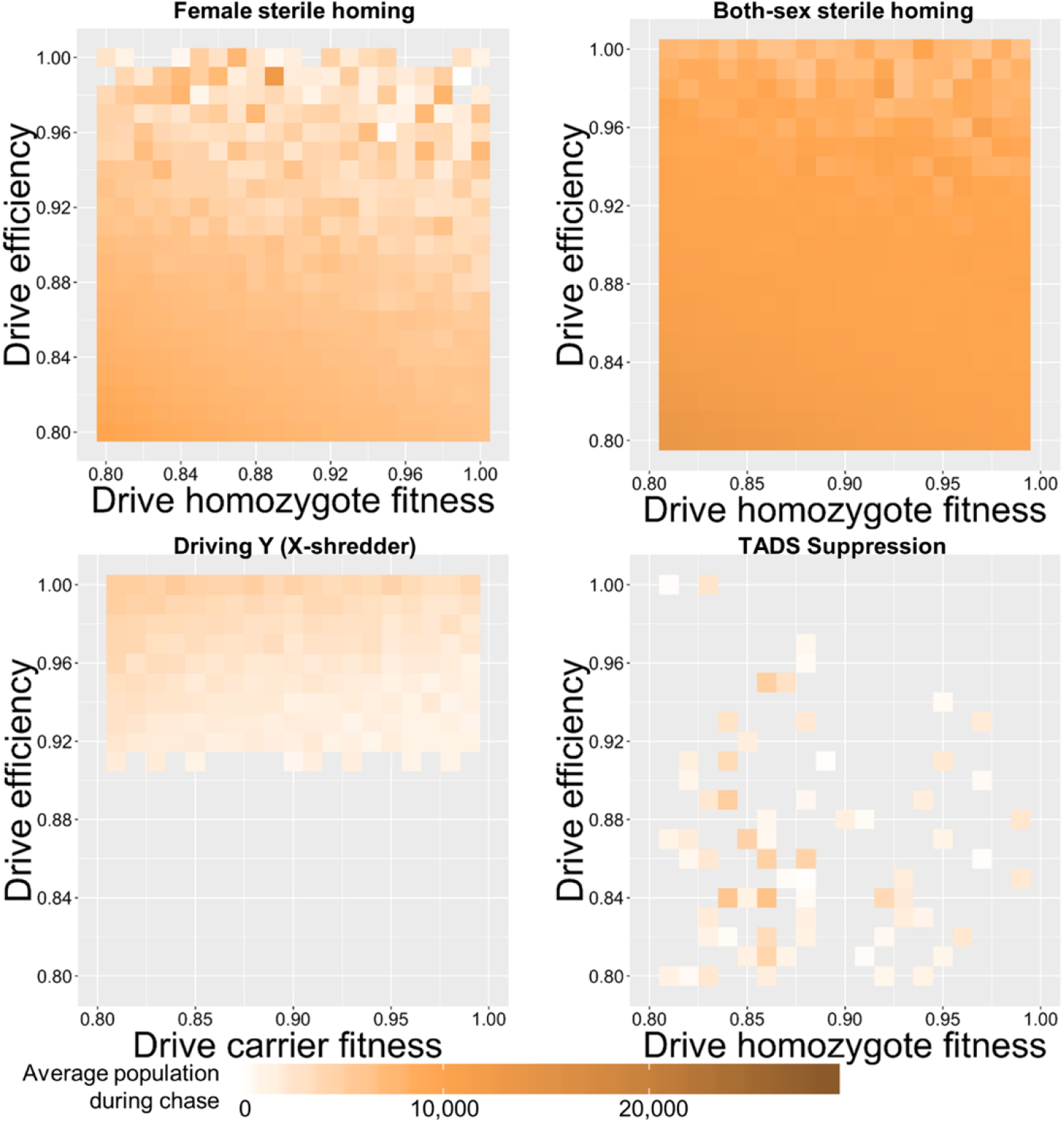
The effect of drive fitness and efficiency on the average population during chasing in continuous space. Drive-carrying individuals were released into the middle of a wild-type population in a 1×1 square. The average population during a chase is shown as the drive fitness and the drive efficiency varies. Each point represents the average of at least 20 simulations, but only simulations that contained a chase were included. Grey areas represent parameter sets in which chasing did not occur.

**Figure S12.**
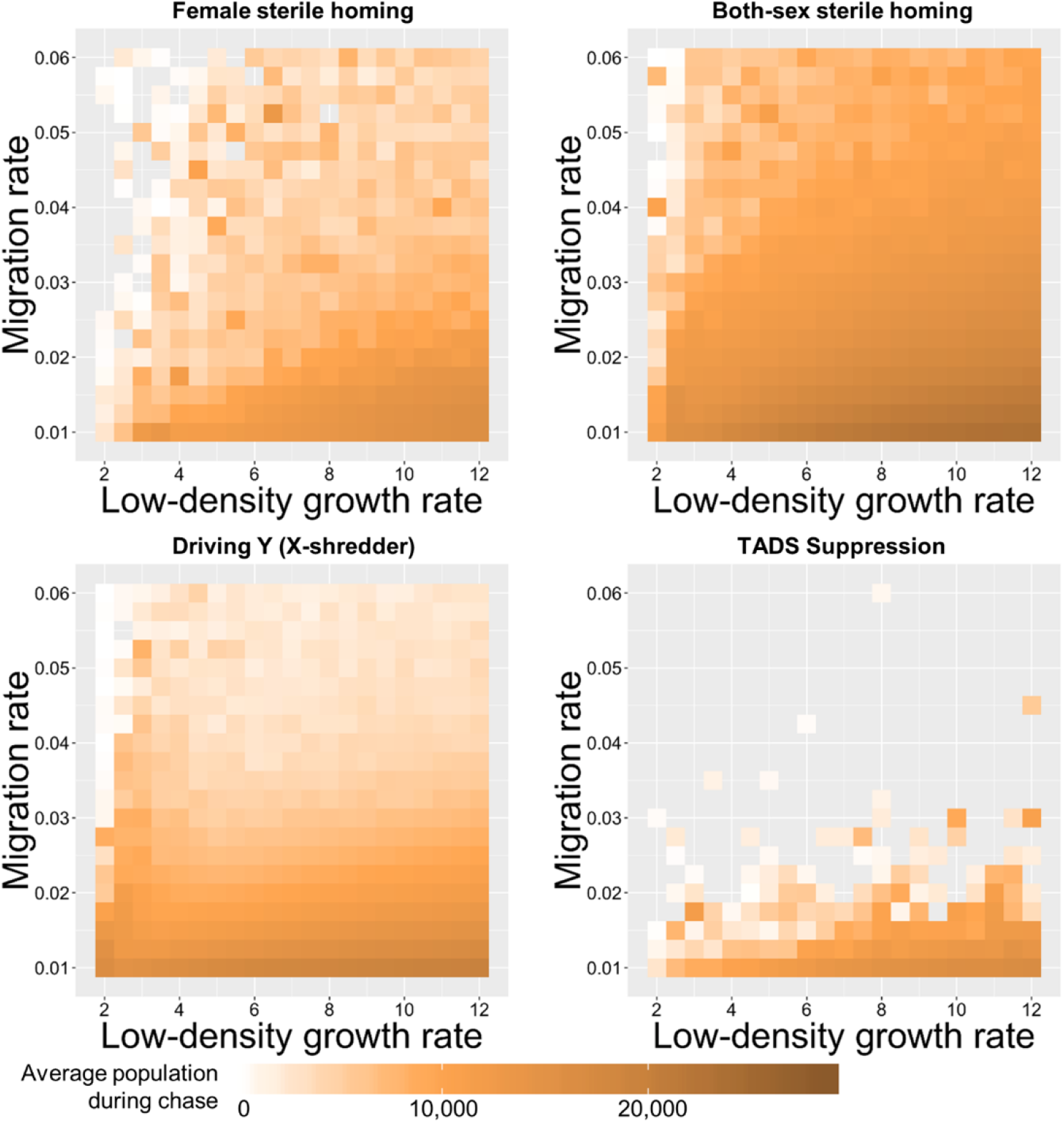
The effect of the low-density growth rate and the migration rate on the average population during chasing in continuous space. Drive-carrying individuals were released into the middle of a wild-type population in a 1×1 square. The average population during a chase is shown as the drive fitness and the drive efficiency varies. Each point represents the average of at least 20 simulations, but only simulations that contained a chase were included. Grey areas represent parameter sets in which chasing did not occur.

### Impact of chasing on spatial distribution of individuals

To further characterize chasing, we examined the average Green’s coefficient for wild-type individuals during a chase, which provides a measure of clustering, with higher values representing greater levels of clustering (Figure 2). We used Green’s coefficient only for wild-type individuals, since this was found to give a greater dynamic range. We found that higher Green’s coefficient was associated with a smaller number of groups of wild-type individuals experiencing chasing, though the size of such groups was more prone to fluctuations when Green’s coefficient was high (Figure 2). When chasing was more common, we found that the average Green’s coefficient during chasing was lower (Figures S13–14), which represented multiple clusters of wild-type individuals experiencing chasing at any given time.

**Figure S13.**
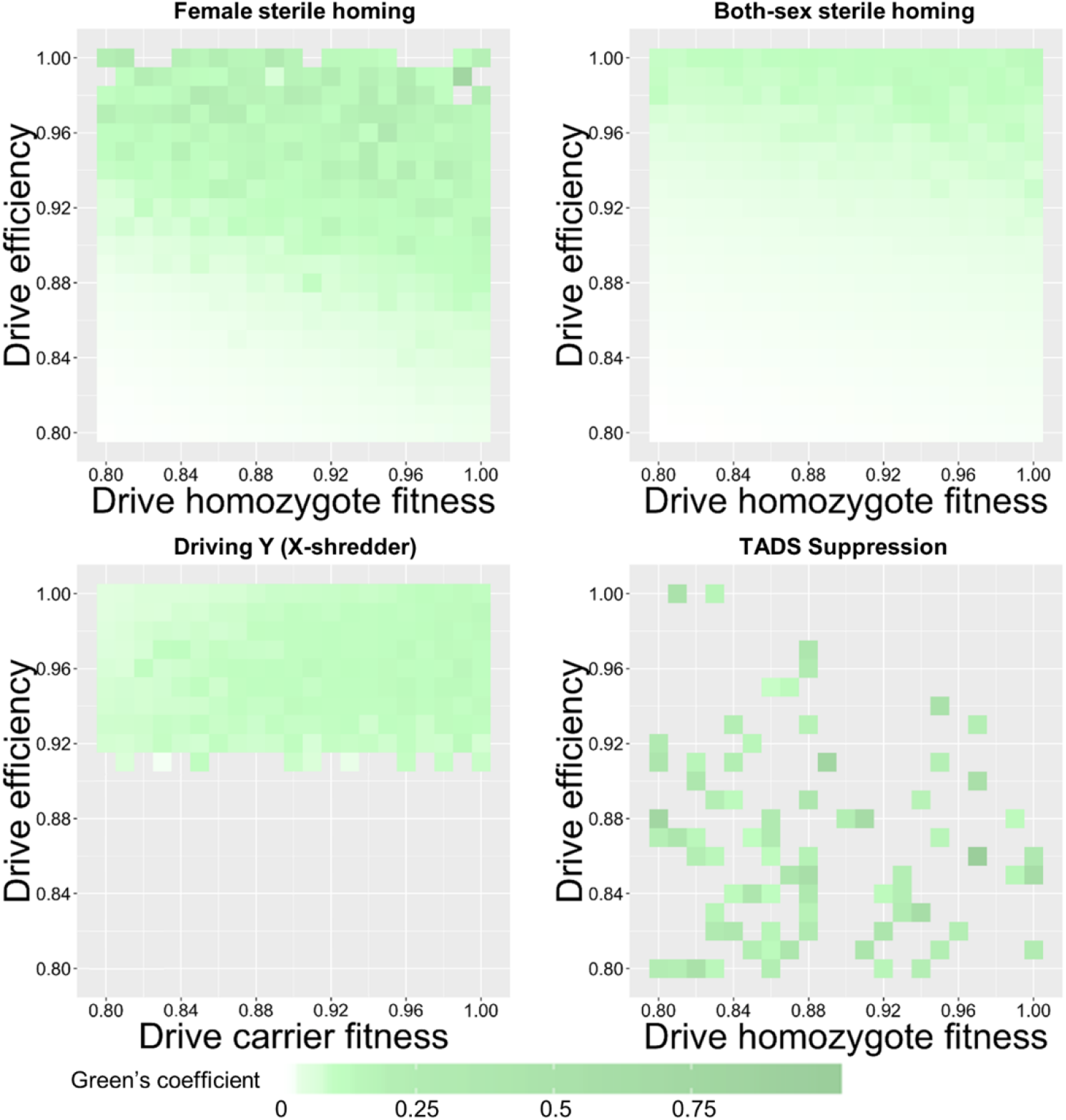
The effect of drive fitness and efficiency on Green’s coefficient during chasing in continuous space. Drive-carrying individuals were released into the middle of a wild-type population in a 1×1 square. Each point represents the average of at least 20 simulations, but only simulations that contained a chase were included. Grey areas represent parameter sets in which chasing did not occur.

**Figure S14.**
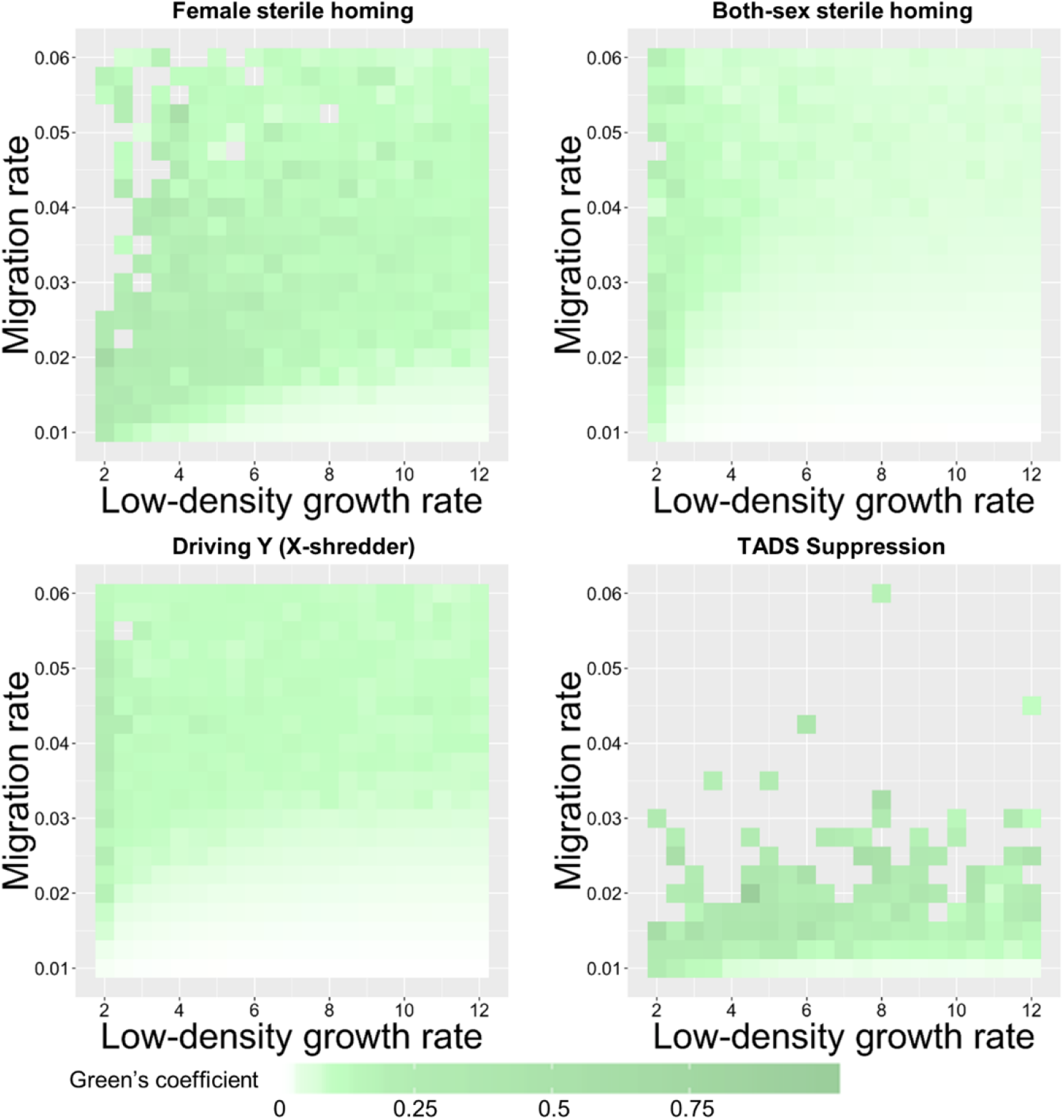
The effect of the low-density growth rate and the migration rate on Green’s coefficient during chasing in continuous space. Drive-carrying individuals were released into the middle of a wild-type population in a 1×1 square. Each point represents the average of at least 20 simulations, but only simulations that contained a chase were included. Grey areas represent parameter sets in which chasing did not occur.

**Figure S15.**
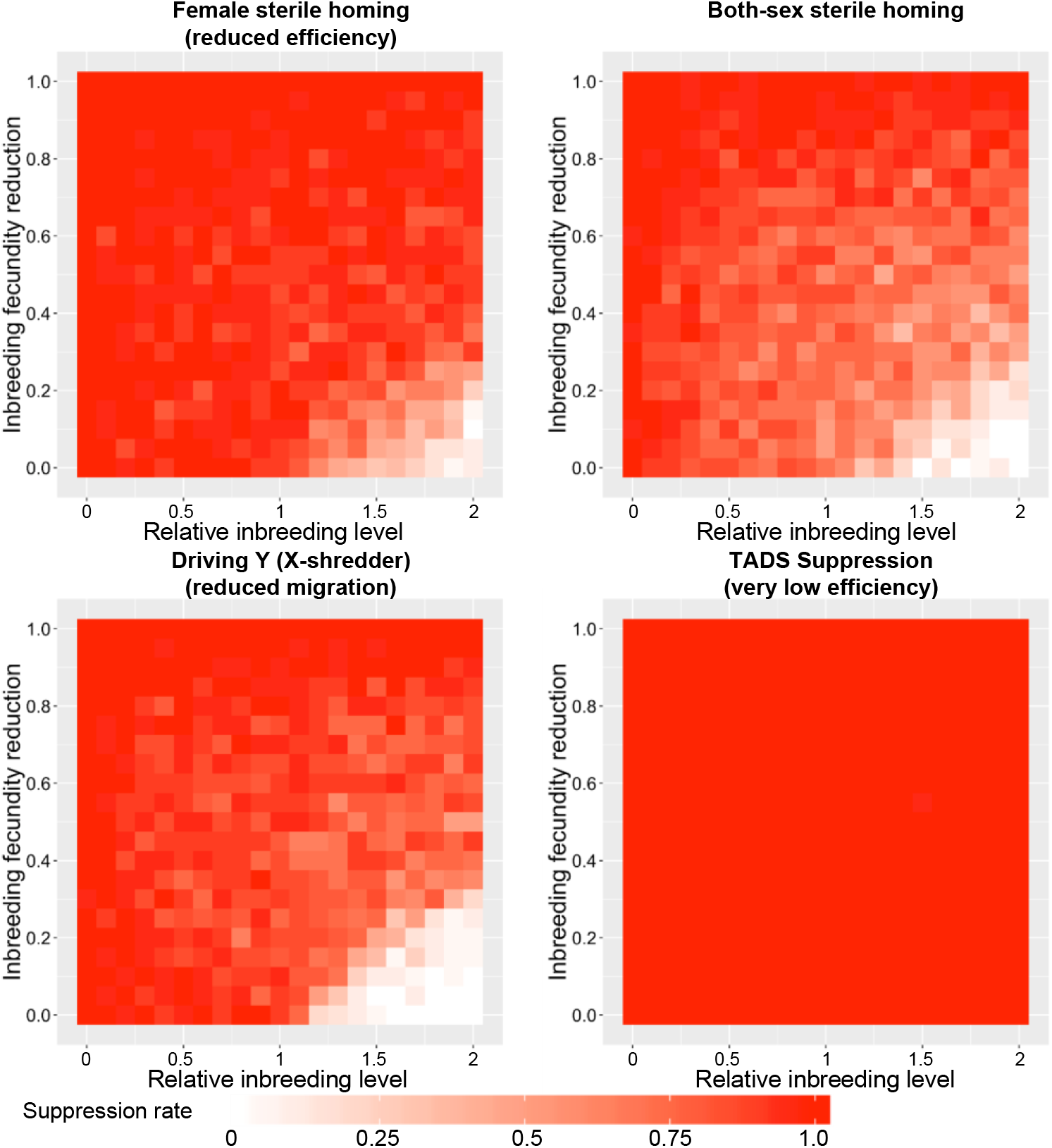
The effect of inbreeding on the suppression rate in continuous space. Drive-carrying individuals were released into the middle of a wild-type population in a 1×1 square. The relative inbreeding level specifies the relative preference a female gives to siblings when choosing a mate as compared to non-siblings (a value of 1 means that no preference is given, while a value of 2 means that siblings are twice as likely to be chosen), before adjustment by fitness. If a female mates with a sibling, she will suffer an inbreeding fecundity penalty (0 = no fecundity reduction for sibling-mated females, 1 = no offspring are generated by sibling-mated females). Each point represents the average of at least 20 simulations. To show a greater dynamic range of outcomes, some default parameters have been modified (female sterile homing drive efficiency and fitness has been reduced to 0.92, the migration rate has been reduced to 0.035, and the low-density growth rate has been increased to 8; the driving Y migration rate has been reduced to 0.0325; TADS suppression drive efficiency and fitness has been reduced to 0.8, the migration rate has been reduced to 0.02, and the low-density growth rate has been increased to 12).

**Figure S16.**
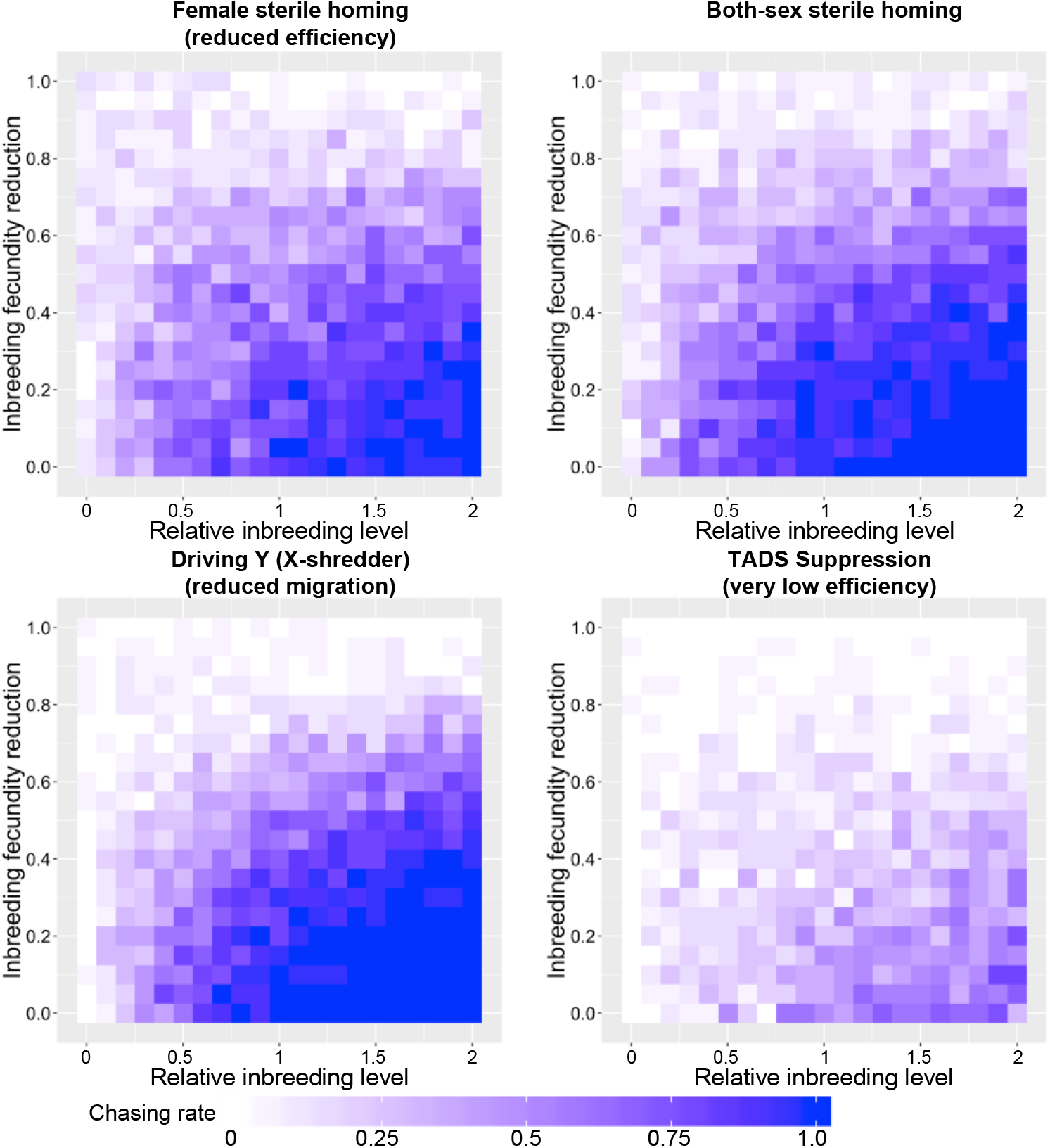
The effect of inbreeding on the rate of chasing in continuous space. Same as Figure S15, but chasing rate is shown instead of suppression rate.

### Initiation of chasing

Chasing appears to start when some wild-type individuals avoid the expanding wave of the drive and move into empty space, giving them an advantage due to reduced competition (Figure 2). To better understand the initiation of chasing, we analyzed a model in which individuals were allowed to move over only one dimension. This model considers the landscape as a one-dimensional line, rather than a square area. Values that represent radii in the two-dimensional model now represent distances along the *x*-axis from the focal point, and areas for determining density are now one-dimensional intervals. Coordinates for individuals are no longer (x,y) pairs, but are only x-coordinates. We used a line of length 5 with *K* = 3,925 to produce the same carrying density (*ρ*_*k*_) as in our two-dimensional model. Finally, the drive is released from the left side of the landscape in an interval of size 0.01, instead of in a central circle. In our one-dimensional model, chasing detection is simplified to determining when, or if, there is a mostly empty population interval with populated areas to its left. Since the drive is always released from the left side of the landscape, this detection indicates that wild-type alleles were able to penetrate the right-moving wave of drive individuals and recolonize empty areas the wave had already passed.

We found that even with one dimension, suppression did not always occur (Figures S17–18), though patterns for all drives excepts TADS suppression were somewhat different than the 2D model. This is partially explained by the greater rate of drive loss in 1D (Figures S19–20), which likely occurs due to the smaller numbers of drive individuals at any time in the advancing 1D wave front of the drive compared to 2D wave fronts. In many cases, loss of the drive in 1D may correspond to loss of the drive in a small patch of the 2D model, allowing wild-type individuals to move through the newly formed “hole” in the advancing drive wave front, initiating chasing by recolonizing empty patches. However, chasing does not require a complete hole in the drive wave front to form. Indeed, in our 1D models, we found generally higher rates of chasing than in 2D (Figures S21–22). In these cases, wild-type individuals must have been able to permeate the drive wave to move into the empty space the drive had just cleared, thus initializing a chase.

**Figure S17.**
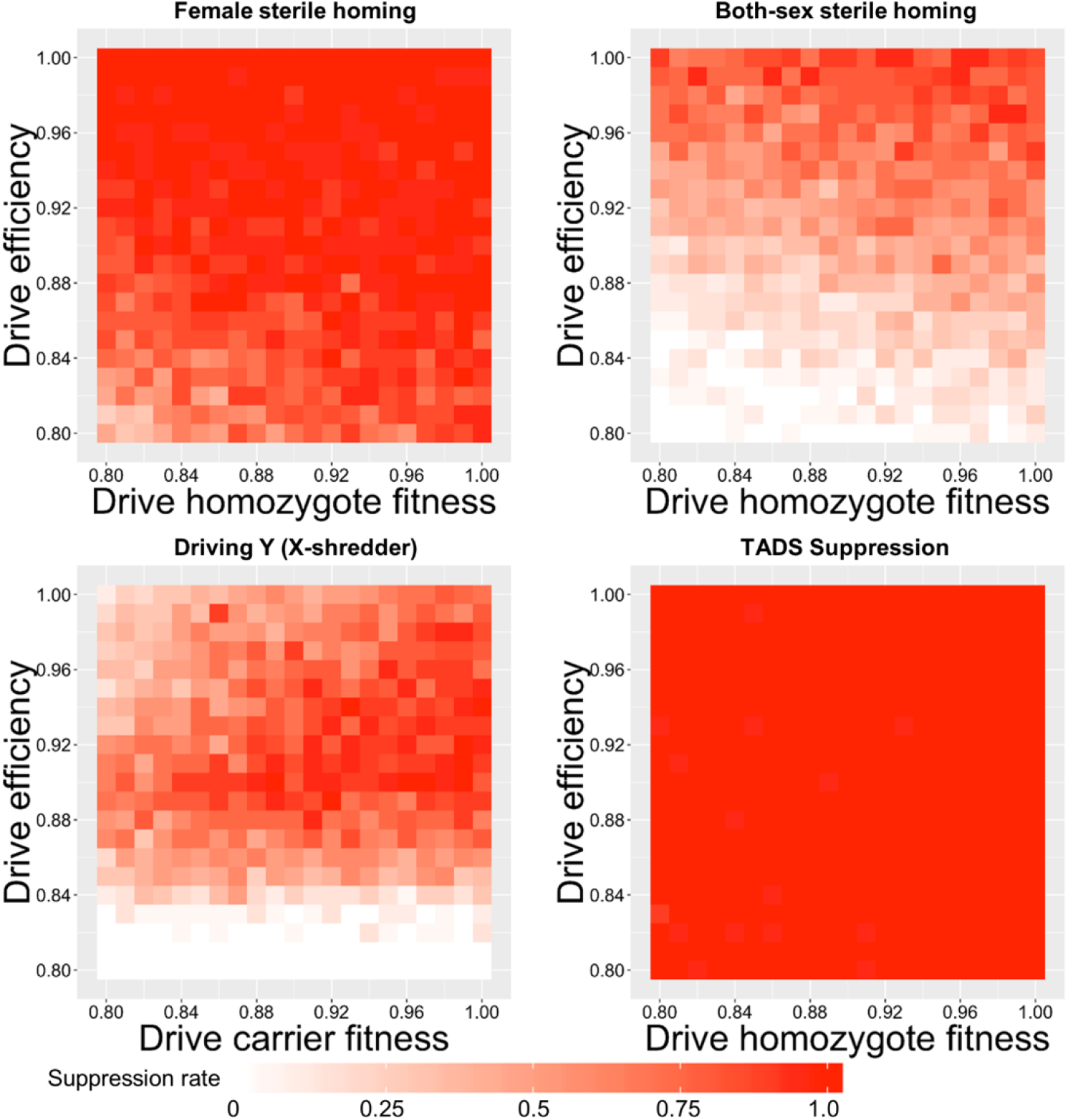
The effect of drive fitness and efficiency on the suppression rate in one-dimensional continuous space. Drive-carrying individuals were released into the left side of a wild-type population in the one-dimensional model. The proportion of simulations where the population was eliminated is shown. Each point represents the average of at least 20 simulations.

**Figure S18.**
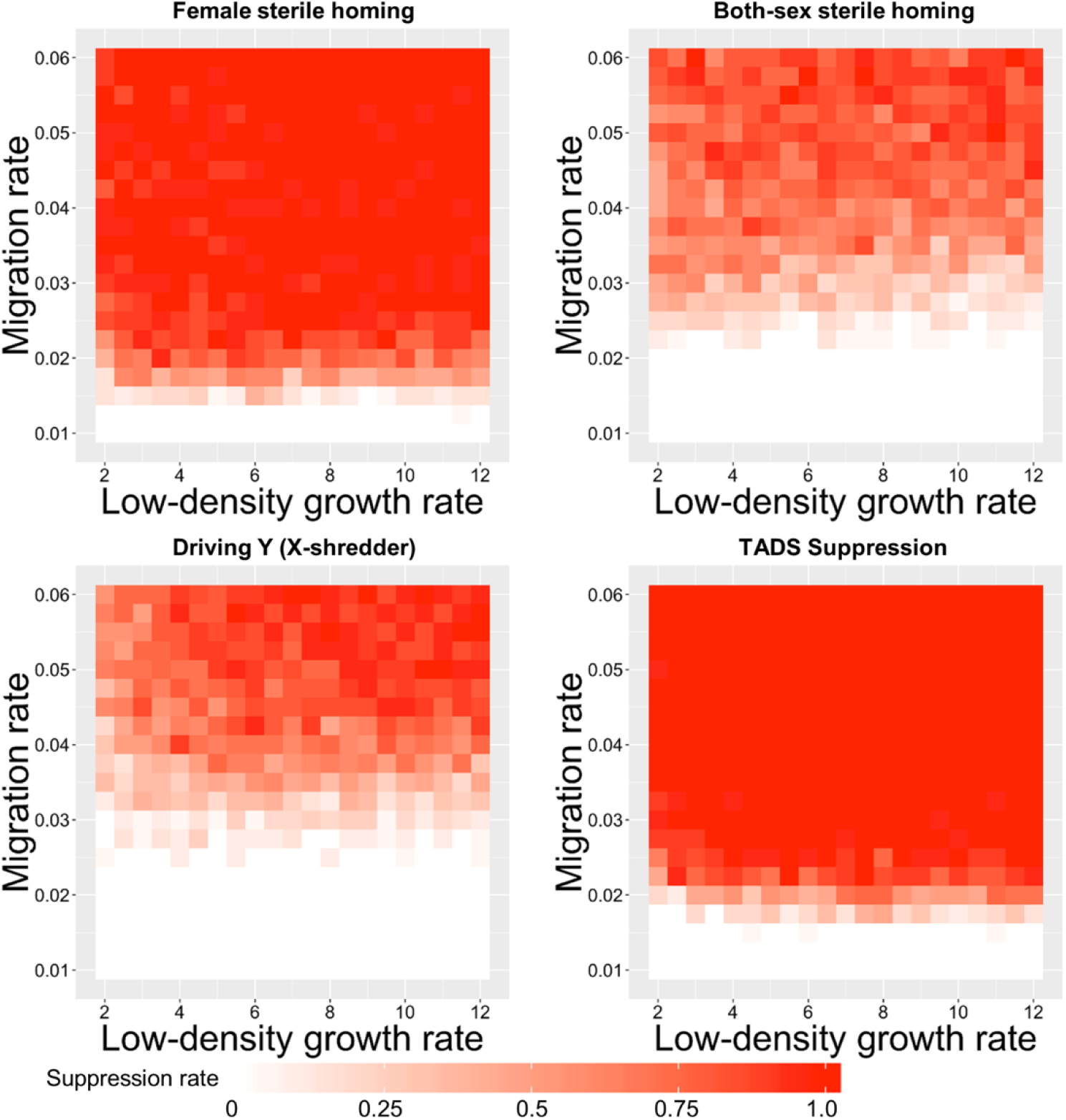
The effect of the low-density growth rate and the migration rate on the suppression rate in one-dimensional continuous space. Drive-carrying individuals were released into the left side of a wild-type population in the one-dimensional model. The proportion of simulations where the population was eliminated is shown. Each point represents the average of at least 20 simulations.

**Figure S19.**
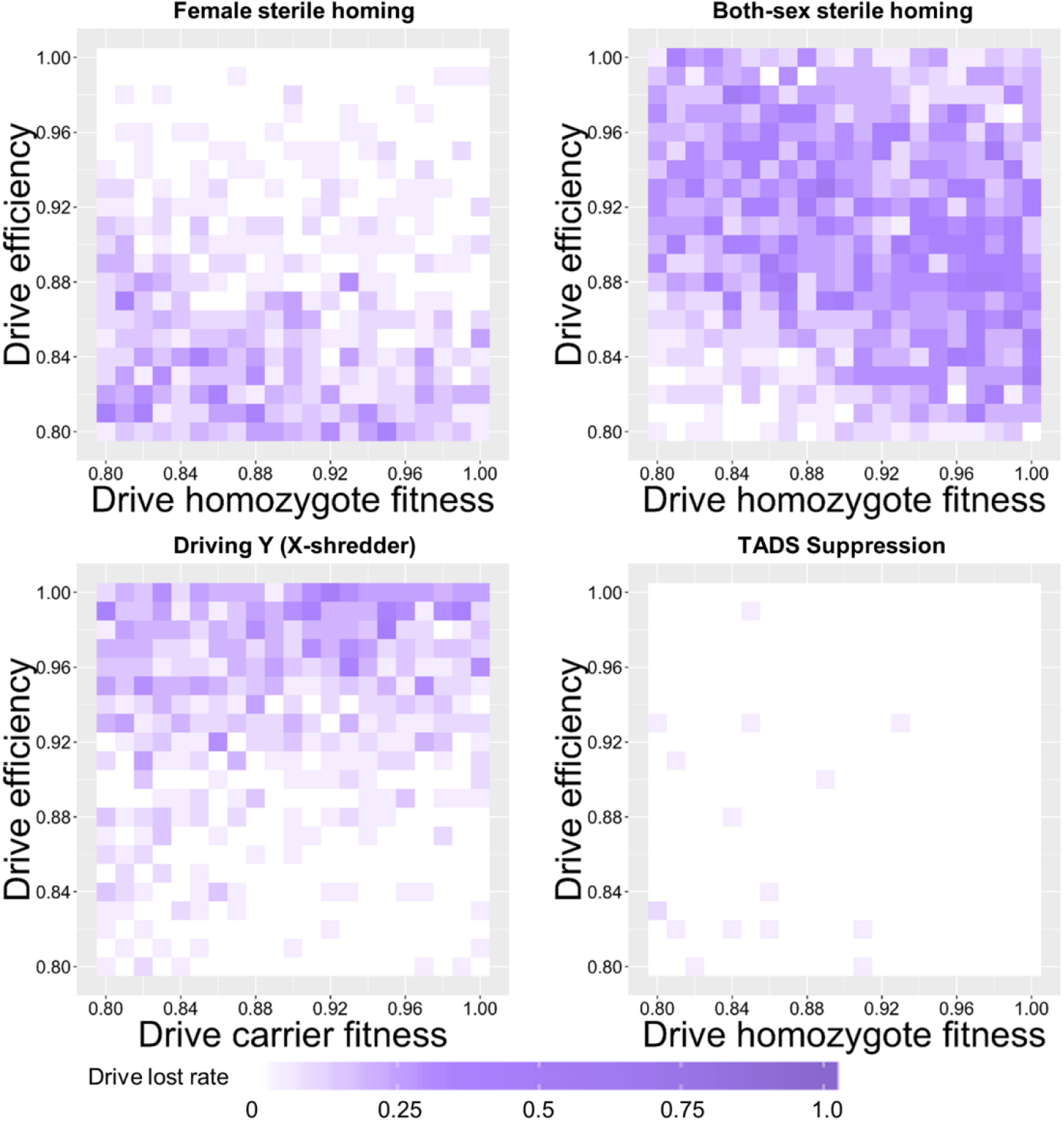
The effect of drive fitness and efficiency on the drive elimination rate in one-dimensional continuous space. Drive-carrying individuals were released into the left side of a wild-type population in the one-dimensional model. The proportion of simulations where the drive was lost is shown. Each point represents the average of at least 20 simulations.

**Figure S20.**
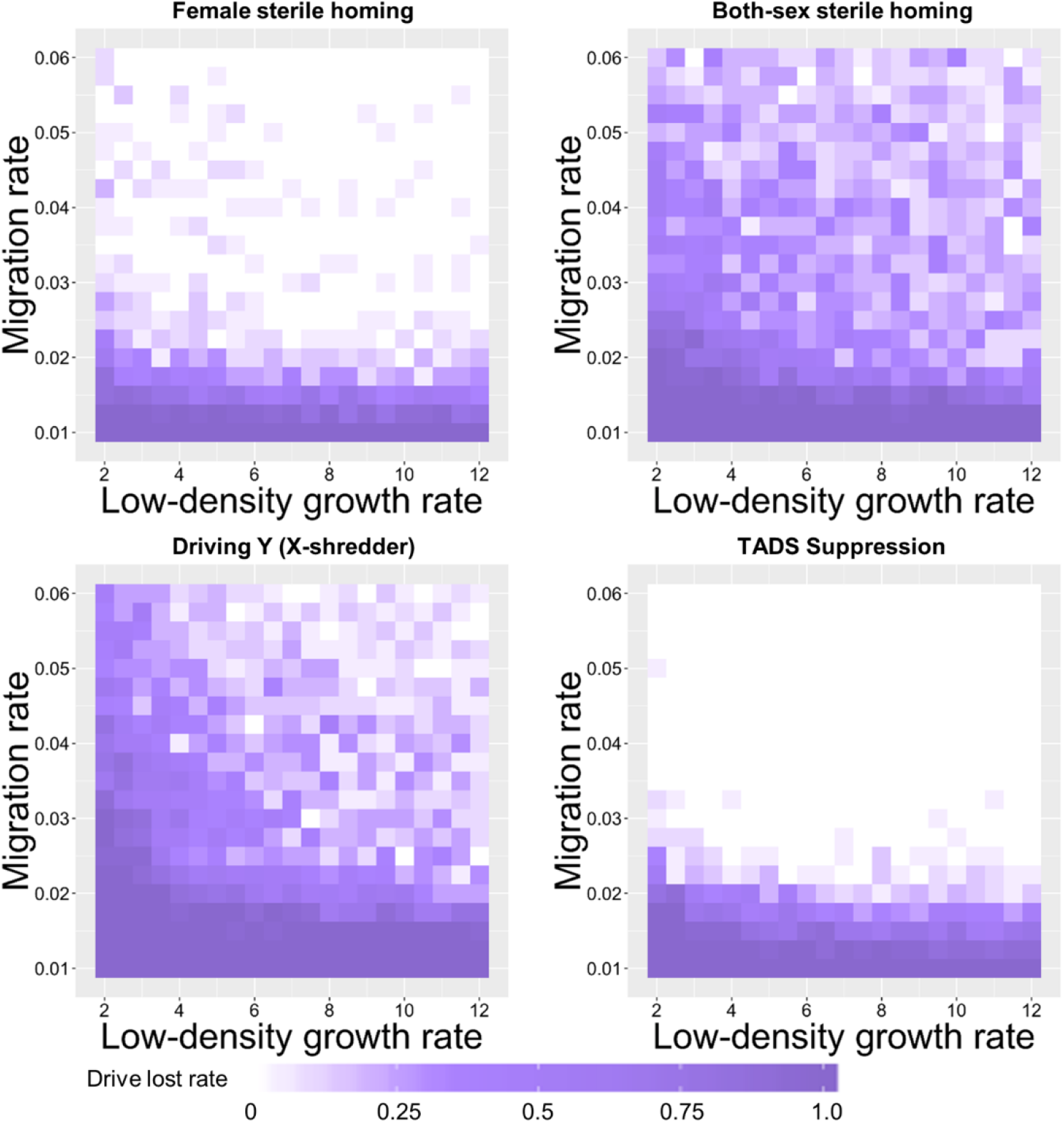
The effect of the low-density growth rate and the migration rate on the drive elimination rate in one-dimensional continuous space. Drive-carrying individuals were released into the left side of a wild-type population in the one-dimensional model. The proportion of simulations where the drive was lost is shown. Each point represents the average of at least 20 simulations.

**Figure S21.**
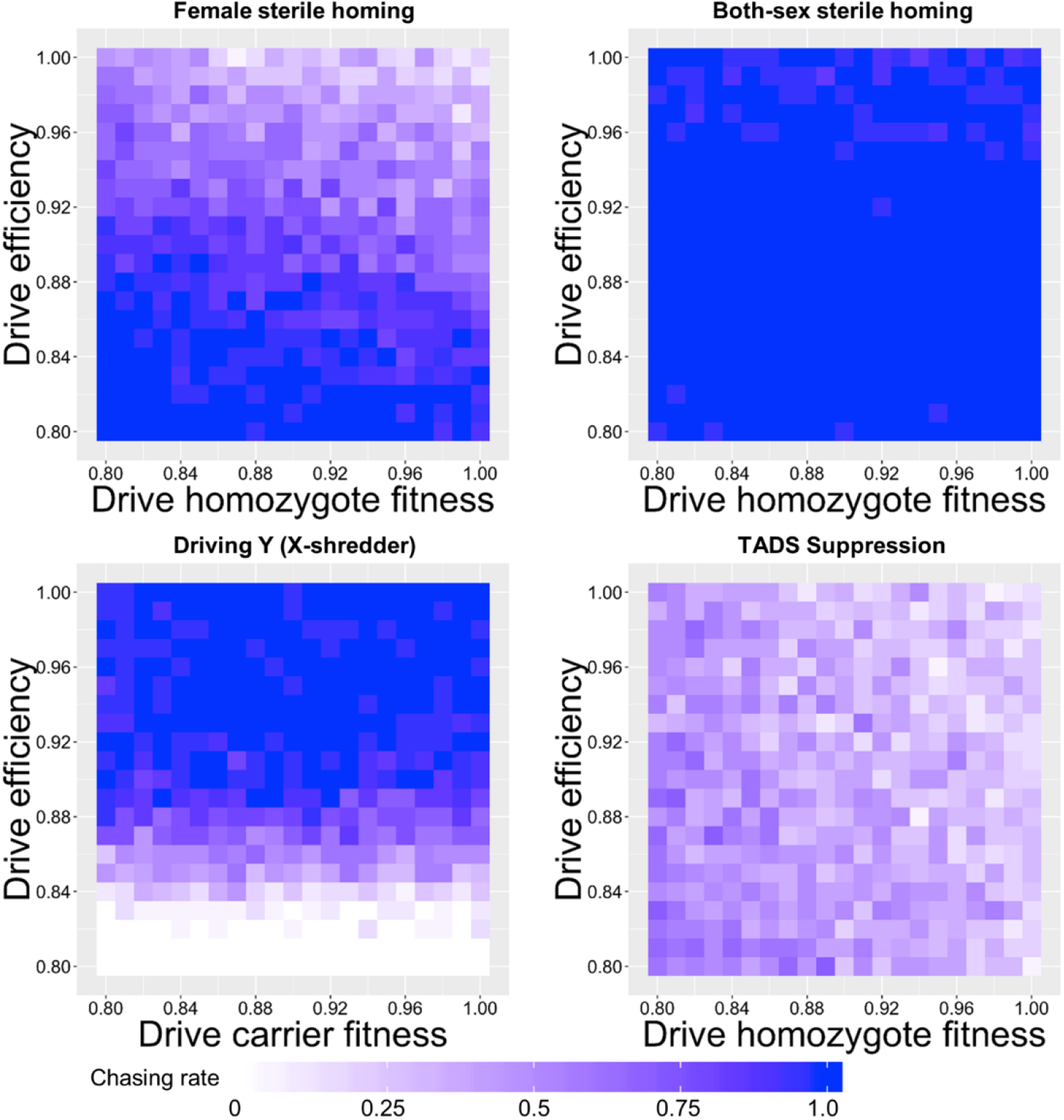
The effect of drive fitness and efficiency on the rate of chasing in one-dimensional continuous space. Drive-carrying individuals were released into the left side of a wild-type population in the one-dimensional model. The proportion of simulations where a chase occurred is shown. Each point represents the average of at least 20 simulations.

**Figure S22.**
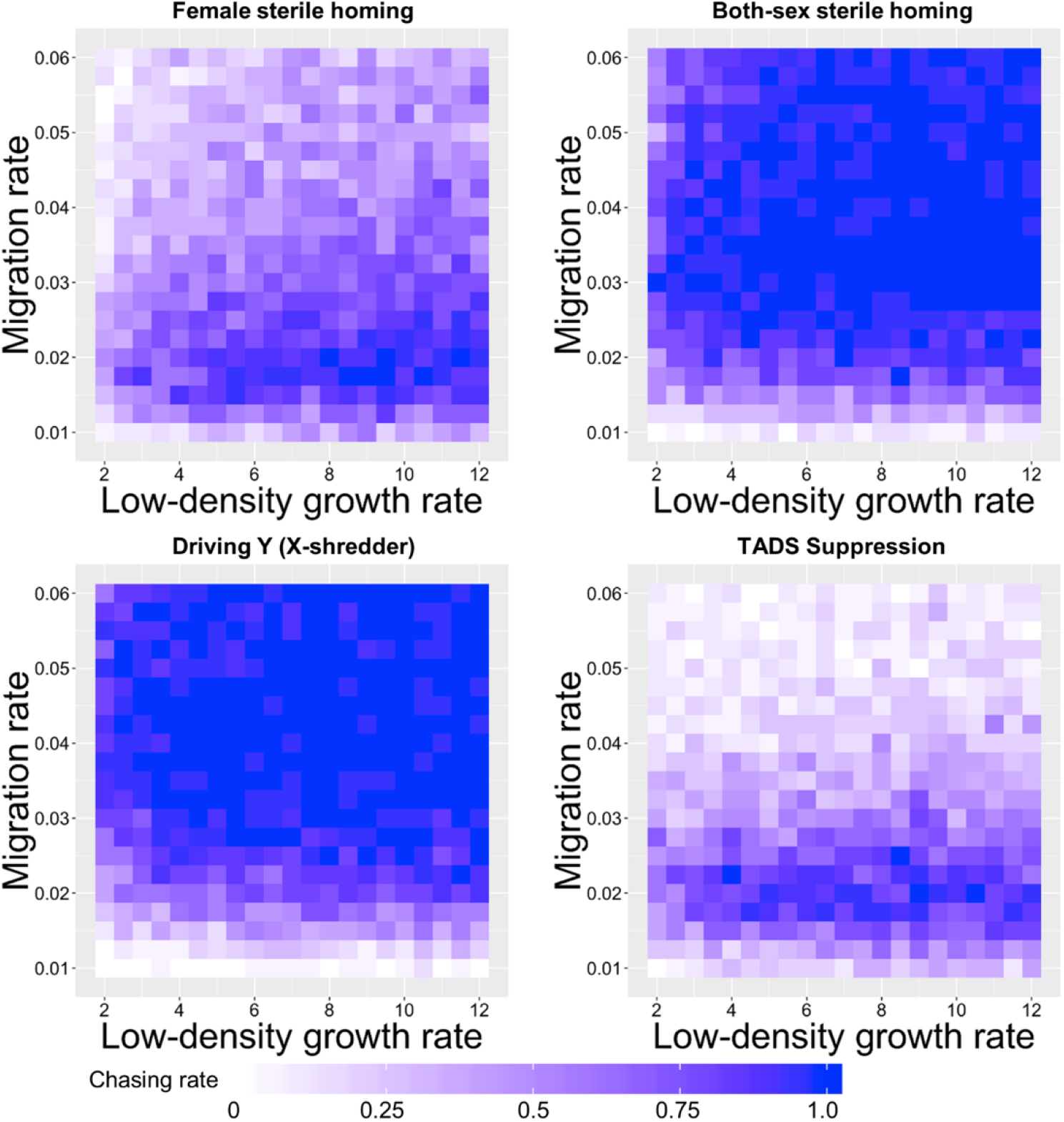
The effect of the low-density growth rate and the migration rate on the rate of chasing in one-dimensional continuous space. Drive-carrying individuals were released into the left side of a wild-type population in the one-dimensional model. The proportion of simulations where a chase occurred is shown. Each point represents the average of at least 20 simulations.

### Comparative analysis of drive types

Our analysis of the data in Figure 3 revealed major differences in the ability of the four drive types to avoid chasing and suppress the population in our spatial model. As an attempt to understand what may be causing these differences, we explored the characteristic differences in how these four drives achieve elimination in the panmictic model. In particular, we studied the population characteristics at the time point when the drive had reached a frequency of 90% in the population (Table S1). This specifies a point where chasing often started in the spatial model. We hypothesized that stochastic effects during this phase may play a key role in determining whether the drive achieves elimination in the spatial model, is lost, or chasing starts. High levels of stochasticity during this phase may allow for the escape of a few wild-type individuals into empty space to start a chase. At the same time, high stochasticity could also support sudden loss of the drive as well as population elimination.

To assess the level of stochastic effects in the panmictic model when the drive reached 90% frequency for each of the four drive types, we first determined the average number of females present in this generation that will actually reproduce (Table S1). For the driving Y, we only included the females that mate with wild-type males, since only these produce significant numbers of females that will determine the reproductive capacity for the next generation. This number is likely more representative of the degree of stochasticity than the total population size, as it is these pairings that generate individuals in the next generation. TADS suppression had an average of 934 pairings producing females, far higher than all the other drives, which correlates with the success of this drive. The female sterile homing driving similarly had far more pairings producing females (114) than the both-sex sterile homing drive (12) and was far less prone to both chasing and drive loss. The Driving Y had an average of 103 pairings producing females, which was closer to the female-sterile homing drive. However, its performance was closer to the both-sex sterile homing drive. Thus, other factors likely account for the relative performance of the Driving Y, which uses a substantially different mechanism of suppression (sex ratio vs. sterility).

**Table S1.** Population characteristics at 90% drive frequency in panmictic scenarios correlate with tendency of drives to initiate chasing.

|  | Female sterile homing | Both-sex sterile homing | Driving Y (X-shredder) | TADS Suppression |
| --- | --- | --- | --- | --- |
| Population size | 1363 | 865 | 10969 | 10999 |
| Fertile females | 7.9% | 8.1% | 9.1% | 50% |
| Sterile females | 43% | 41% | N/A | N/A |
| Fertile males | 49% | 8.8% | 8.2% | 8.3% |
| Sterile males | N/A | 42% | 83%* | 42% |
| **Expected number of reproducing females** | **114** | **12** | **103\*** | **934** |
| **Chasing rate** | **medium** | **very high** | **high** | **low** |

Values are the average over 20 simulations
\*Drive-carrying males for the Driving Y
\*\*For the Driving Y, this includes only females expected to mate with wild-type males

Another way to compare the drives is to consider the distributions of genotype frequencies in the advancing drive wave in our one-dimensional model (Figure S23, see section “Initiation of chasing” for a definition of this model). This facilitates a qualitative assessment of how easy it might be for wild-type individuals (or alleles) to permeate the advancing drive wave and reach empty space behind it, thus starting a chase. We found that the female-sterile drive has relatively high numbers of drive individuals between the leftmost wild-type individuals and empty space. This indicates that wild-type individuals would have a more difficult time reaching this empty space. The both-sex sterile homing drive has lower numbers of drive individuals in this region. For the Driving Y, comparisons are more difficult, due to the markedly different genotypes. However, wild-type females are always needed to reproduce and are found in roughly the same quantities as wild-type males in any given area. Since females are required for production, the leftmost females may occasionally mate with a nearby wild-type male and distribute their offspring to the left of new drive males, thus allowing for initiation of a chase. Finally, the TADS suppression drive has very high numbers of drive individuals between empty space and the left-most point where wild-type individuals are present. While TADS suppression drive may advance somewhat more slowly than the other drives, this may allow it to largely avoid chasing behavior.

During a chasing situation, the ability of the wild-type individuals to avoid a drive and continue chasing is related to the relative rates of advance of the wild-type and drive. We thus determined the difference between the rate of advance of wild-type individuals into empty space and of the drive into wild-type populations (Figures S24–25). Usually, wild-type advanced faster into empty space than the drive advanced into a wild-type population. This is likely a prerequisite for robust chasing and was the case regardless of drive fitness or efficiency, though less effective drives did advance slower. An increase in the low-density growth rate increased the relative rate of wild-type advance, while increased migration rate tended to reduce the difference between wild-type and drive. When the migration rate or the low-density growth rate was very low, however, the drive was able to advance faster than wild-type individuals. Chasing in this parameter space is likely sustained by stochastic factors, which are increased under these conditions.

**Figure S23.**
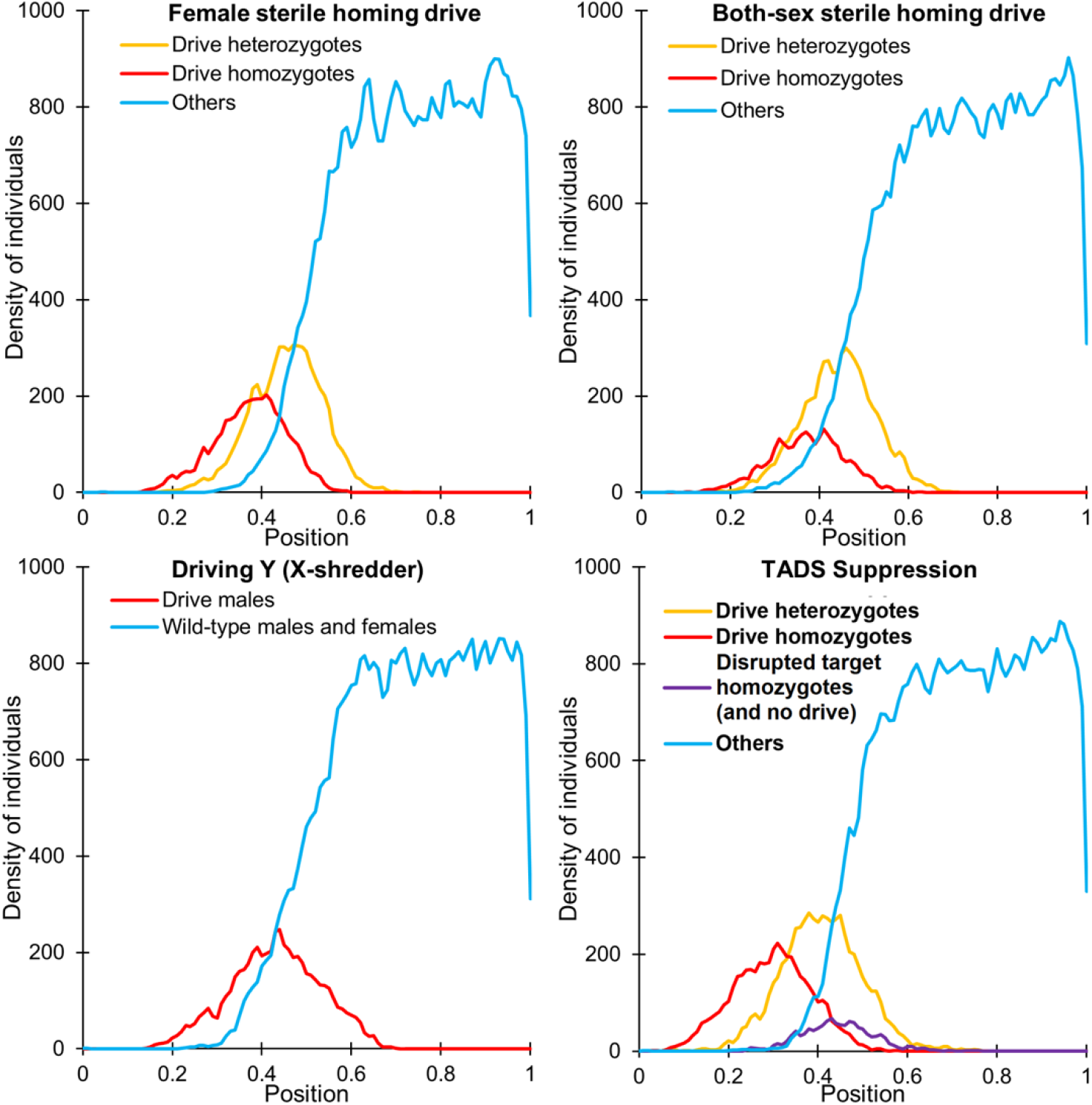
Frequency distribution of genotypes in an advancing drive wave. 800 drive heterozygotes (drive-carrying males for the Driving Y) were released between coordinates 0 and 0.01 into a population of 80000 wild-type individuals in a one-dimensional space of length 1. The genotype and position of each individual were tracked for each generation. Typical genotype distributions at a particular generation once the wave of advance had formed are shown. To make differences between drives clearer, the migration rate has been reduced from its default of 0.04 to 0.03. Density of individuals is the number within the density interaction radius of the position. A higher carrying density than our default value was used to minimize the impact stochastic fluctuations. Nearly all individuals in the “others” category are homozygous for wild-type alleles.

**Figure S24.**
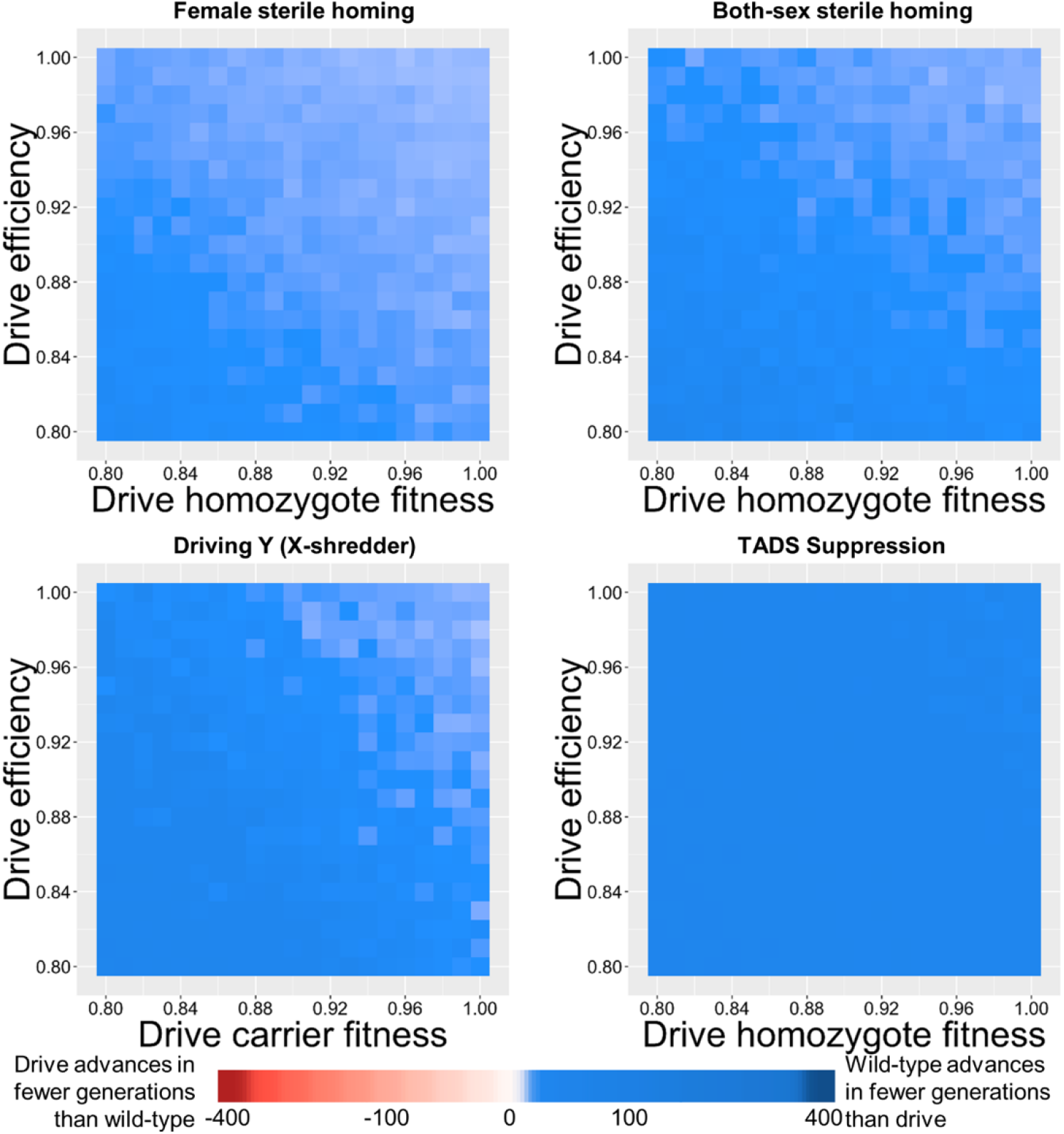
The effect of drive fitness and efficiency on drive and wild-type wave propagation rates in one-dimensional continuous space. Drive-carrying individuals were released into the left side of a wild-type population in the one-dimensional model. Additionally, 39 wild-type individuals were released in a similar but empty space. The difference in the number of generations to advance a distance of half the arena length between the drive (advancing into a wild-type population) and the wild-type (advancing into an empty space) is shown. Each point represents the average of at least 20 simulations.

**Figure S25.**
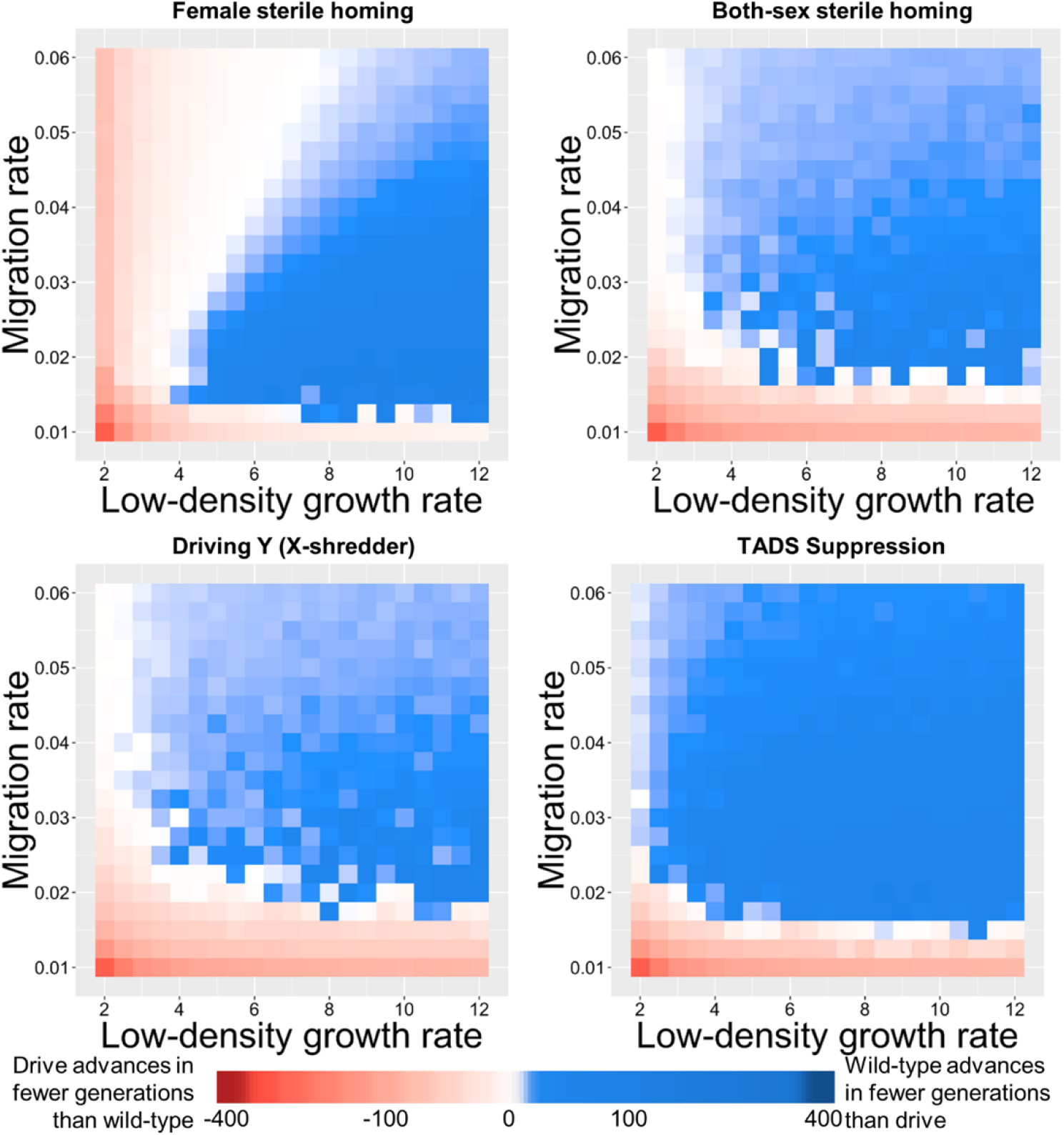
The effect of the low-density growth rate and the migration rate on drive and wild-type wave propagation rates in one-dimensional continuous space. Drive-carrying individuals were released into the left side of a wild-type population in the one-dimensional model. Additionally, 39 wild-type individuals were released in a similar but empty space. The difference in the number of generations to advance a distance of half the arena length between the drive (advancing into a wild-type population) and the wild-type (advancing into an empty space) is shown. Each point represents the average of at least 20 simulations.

